# Anaesthesia selectively decouples signal propagation from responsivity in the fronto-parietal loop

**DOI:** 10.1101/2025.04.15.648923

**Authors:** Abhilash Dwarakanath, Majid Khalili-Ardali, Marion Gay, HV Raghuram, Maxime Roustan, Bechir Jarraya, Theofanis I Panagiotaropoulos

## Abstract

Loss of consciousness under anaesthesia is accompanied by widespread silencing of neurons, and disruption of cortical dynamics. Yet, how this affects mesoscale signal propagation within higher-order associative areas, crucially implicated in theories of consciousness, remains poorly understood. Here we combined intracortical microstimulation, and simultaneous multielectrode recordings in the ventrolateral prefrontal cortex (vlPFC) and the posterior parietal cortex (PPC) of macaques across wakefulness, and graded depths of anaesthesia. Spiking responses revealed distinct regimes: in the PFC, higher-amplitude stimulation elicited a delayed single rebound after sustained inhibition, whereas in the PPC, a faster and double-rebound profile emerged. Despite enhanced local spiking and LFP responsivity under anaesthesia, we found a strong and selective suppression of lateral signal propagation in the PFC - a breakdown strikingly absent in the PPC. This dissociation suggests that anaesthesia disrupts consciousness not merely by silencing cortical populations, but by impairing mesoscale integrative processes critical for neuronal dynamics at multiple scales. Our findings demonstrate in a causal and spatially-resolved manner, that lateral signal spread within higher-order cortex is a key mechanism underlying conscious awareness, and its loss under anaesthesia.

## Introduction

Consciousness, a fundamental aspect of human experience, encompasses a spectrum ranging from full awareness and altered states of consciousness to different levels of unconsciousness seen in anaesthesia and consciousness disorders^1–4^. Understanding the neural correlates of such unconscious states, and the disruptive effects of these states on normal cortical functions, has significant implications for elucidating the cortical mechanisms underlying conscious awareness and for the treatment of consciousness disorders^5–7^.

Multiple theoretical frameworks have been developed to address how consciousness arises in the brain^8,9^. Among theories that ascribe a central role to higher cortical and association areas^10–14^, the Global Neuronal Workspace Theory (GNWT) proposes that conscious experience arises from a higher-order, cortex-mediated ignition, amplification and widespread dissemination of sensory information across the brain, with the prefrontal cortex (PFC) playing a key role in this global broadcasting process^15–18^. In contrast, Integrated Information Theory (IIT) posits that consciousness corresponds to the amount and structure of integrated information (*phi* - Φ) generated by a system, emphasising locally maximised, irreducible cause-effect structures within highly interconnected neuronal ensembles, often associated with a posterior cortical “hot zone” ^1,19–21^, rather than with any specific role for these higher order associational cortical areas. Higher-order thought (HOT) ^22,23^ and related theories likewise emphasise the contribution of higher cortical regions to conscious access, albeit with distinct commitments to representation, state, access, hierarchy, and mechanism/dynamics, while recurrent processing and posterior hot zone accounts argue that posterior cortices ^24–26^ may be sufficient for conscious experience, with PFC contributions confined to report, metacognition or task demands ^27^.

Studies in humans using non-invasive whole-brain recording techniques such as BOLD-fMRI have implicated the PFC during conscious, accessible perception of a stimulus^28–30^. More specifically, direct electrophysiological recordings in the PFC show that prefrontal neuronal populations represent the contents of consciousness, while ongoing low-frequency fluctuations in the prefrontal state are involved in the stability and update of these contents^31–35^, even as some authors remain sceptical about a necessary role of PFC for all forms of conscious experience ^36,37^, (but see ^11^.

In terms of known cytoarchitecture at the micro- and mesoscales, the PFC is of particular interest due to its dense reciprocal connections with other cortical areas, especially the parietal cortex, forming a fronto-parietal loop (FP-loop)^38–41^, specifically mediated *via* long-range, bidirectional, monosynaptic projections from layer 5 pyramidal (L5p) neurons. Crucially, beyond conscious perception *per se*, this fronto-parietal network is widely believed to be central for sensory awareness and goal-directed access, which are either suppressed, or abolished under anaesthesia and similar states^15,42,43^. Work exploring posterior thalamocortical interactions *via* deep brain stimulation (DBS), and pharmacological manipulation of associational regions – across species – has both highlighted that the precise contribution of fronto-parietal circuitry to consciousness ^44^, and muddied the contribution of posterior cortical regions, which are supported by studies employing gross perturbational and complexity metrics ^45,46^. Therefore, to critically address the role of these areas, in particular the ventrolateral prefrontal cortex (vlPFC) and the posterior parietal cortex (PPC), in the genesis and loss of conscious awareness, direct causal perturbational studies at the micro- and mesoscale are necessary^10,36^.

Studying these regions under anaesthetics, which cause reversible unconsciousness, can help reveal mechanisms of consciousness loss and recovery^47–51^, and can thus inform competing accounts by probing how state-dependent changes in effective connectivity and mesoscale propagation differ across key cortical nodes. Different anaesthetics have been used to understand local and global neural dynamics during conscious and unconscious states^47,52^. For GABAergic anaesthetics, cortical states are dominated by anteriorised slow rhythms^53–57^ in the delta-theta range (1-4 and 5-8 Hz)^58,59^, exhibiting high intra-areal but low inter-areal coherence^55,58,60^. Sevoflurane, a commonly used veterinary inhalant anaesthetic, potentiates inhibitory neurotransmission by primarily enhancing GABA_A_ receptor function^48,61^. At the cellular and microcircuit level, general anaesthesia can decouple cortical pyramidal neurons ^62^ and alter dendritic integration in a way that profoundly reshapes how inputs are transformed into output activity, as shown for example in the somatosensory cortex under ketamine-xylazine ^63^. Furthermore, sevoflurane reconfigures cortical neuronal discharge activity from diffuse and decorrelated to distinct UP/DOWN states, with increased local neuronal correlations^48,64^ while impairing functional connectivity^65–67^. Crucially, sevoflurane disrupts fronto-parietal communication^65,66^ while also leading to a reduction in large-scale signal complexity (as measured by the Perturbational Complexity Index), a signature of the level of consciousness^47,68^. Related work combining recordings and deep brain stimulation has shown that loss and recovery of consciousness are accompanied by characteristic changes in large-scale dynamics, including within fronto-parietal circuits and thalamus ^44,55,69,70^. However, the exact nature of this functional reorganisation at the neuronal population level, in particular at the mesoscale, remains uncharacterised^71,72^. While studies have measured the complexity of large-scale brain activity and distinct states elicited by transcranial magnetic stimulation (TMS) perturbation using scalp EEG^45,73^, investigations specifically targeting nodes of the fronto-parietal loop to characterise the effects of loss of consciousness on mesoscale signal propagation are lacking.

This study aims to address three important questions at the mesoscale cortical level: (1) How does the response profile of prefrontal and parietal populations to electrical stimulation differ across awake and anaesthetised states? (2) How do these states affect signal propagation in time and space? (3) Are there differences in the thus inferred network properties between these two subunits of the FP-loop? To answer these questions, we employed a simultaneous electrical microstimulation-recording protocol during quiet wakefulness (QW), light (LS) and deep anaesthesia (DS), all performed in the same experiment and recording from the same population, to maintain sample consistency. The vlPFC and PPC were recorded in different animals and sessions, a constraint imposed by experimental feasibility that we address in detail in the Discussion. Apart from quasi-similar population response profiles in the two cortical regions, our results reveal a specific signature of anaesthesia in the PFC involving suppression of lateral electrical signal propagation that is absent in the PPC. In contrast, the amplitude of responses following electrical microstimulation did not correlate monotonically with the level of anaesthesia, since deep anaesthesia resulted in responses that were stronger compared to quiet wakefulness.

## Results

We recorded and stimulated from the ventrolateral prefrontal cortex of two male macaques during quiet wakefulness and anaesthesia, in the same experimental sessions, using a 10 x 10 Iridium Oxide coated Utah Array. Specialised adapters allowed us to stimulate and record members of the same population, simultaneously (Fig. 1A).

**Figure 1.**
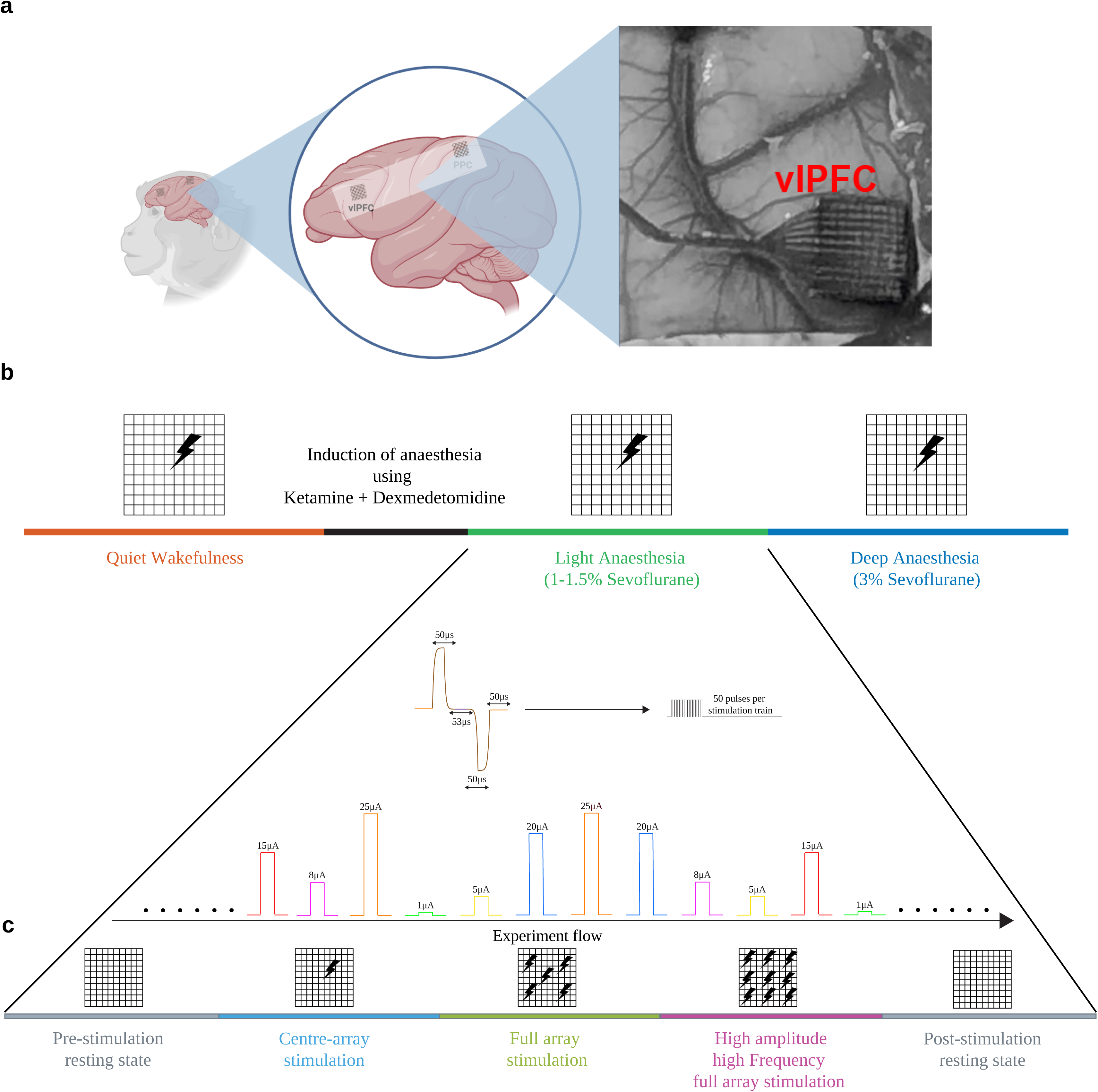
Experimental design and ROI. A. Multielectrode array in the inferior convexity of the PFC, thus targeting the fronto-parietal loop, placed anterior to the bank of the arcuate sulcus and below the principal sulcus, thus being localised to the left ventrolateral prefrontal cortex B. Cortical population activity was recorded in four experimental phases, viz. Quiet wakefulness (QW), transition to anaesthesia and preparation, light anaesthesia (LS - 1-1.5% Sevoflurane), and deep anaesthesia (DS - 3% Sevoflurane), along with the delivery of intracortical microstimulation. The stimulation was devised as a train of 50 biphasic pulses lasting 200ms, with varying amplitudes, presented randomly with an inter-stimulation interval between 10-20s. C. At the start of each experimental epoch, 15-20 mins of resting state activity was recorded, followed by the three stimulation regimes, viz. Centre-array stimulation, full-array stimulation, and full-array high-frequency, high-amplitude stimulation, finally followed by 15-20 mins of post-stimulation resting state.

After recording and electrical microstimulation periods during quiet wakefulness, we induced anaesthesia with a single injection of ketamine and dexmedetomidine. After waiting for ∼90 minutes for the ketamine to metabolise, light anaesthesia was induced by inhalation, and maintained between 1-1.5% of sevoflurane, a GABAergic anaesthetic. After recording and microstimulation during light anaesthesia, we increased the sevoflurane concentration to 3%, inducing deep anaesthesia (Fig. 1B). While most intracortical microstimulation (ICMS) that intend to characterise population response properties employ single-pulse stimulation, we devised our stimulation to mimic those used in studies that seek to, for e.g., augment stimulus detection, or influence behavioural outcome^74–77^. This stimulation, designed as a biphasic 50-pulse train at 250 Hz, was delivered to the two electrodes in the centre of the array, each with opposing polarity, at varying strengths, viz. 1 to 100µA (Fig. 1B). Each stimulation was spaced 10-20s apart to allow sufficient time for cortical activity to return to baseline activity post-stimulation. Resting state periods, both during wakefulness and anaesthesia, were recorded in pre- and post-stimulation periods (Fig. 1C).

### State characterisation in wakefulness and different anaesthesia depths

How does the activity of the same cortical population vary between quiet wakefulness and different levels of anaesthesia?

To address this, in each session from each recorded area in each animal, we selected a population of neurons based on their firing rate during QW (firing rate >= 0.5 spks/s), and used the same population for all downstream analyses. In Monkey A in the PFC, while the firing rates during LS and DS were significantly lower than during QW (p_Comp_ < 0.001), they were not different from each other (p_Comp_ > 0.05). In Monkey C in the PPC however, there were significant differences in the firing rate between QW and DS, and LS and DS (p_Comp_ < 0.001), but not between QW and LS (p_Comp_ > 0.07). In general, the firing was higher during QW, followed by LS and then DS, subject to statistical tests as above. In Monkey B in the PFC, due to very sparse firing, no significant differences were found in the firing rates between any of the 3 conditions (all p_Comp_ > 0.5). The firing rate comparisons are summarised in SIFig. 1. Because each session contained the same number of channels recorded, for this statistical comparison, the Student’s t-test was employed.

Qualitatively, quiet wakefulness is typified by neuronal discharge activity which is diffuse and uncorrelated between channels (Fig. 2A (PFC) and Fig. 2E (PPC) - top row), accompanied by low amplitude high frequency field fluctuations (gamma) (Fig. 2C (PFC) and 2G - top row). The induction of anaesthesia leads to a transition from this decoherent state to periods of coherent discharge interspersed by silence, known as UP/DOWN states, possibly leading to burst suppression in DS (Fig. 2A (PFC) and Fig. 2E (PPC) - middle row (LS), and bottom row (DS))^78,79^. This is reflected in significantly lower delta-theta power, and significantly higher beta and gamma power during wakefulness (p_Comp_ < 0.0001; SIFig. 2), whereas both states of anaesthesia are dominated by low-frequency field activity (Figs. 2D (PFC), 2H - PPC). In both recorded areas, the power in the low-frequencies increased with increasing depth of anaesthesia (LS vs DS; p_Comp_ < 0.0001; SIFig. 2).

**Figure 2.**
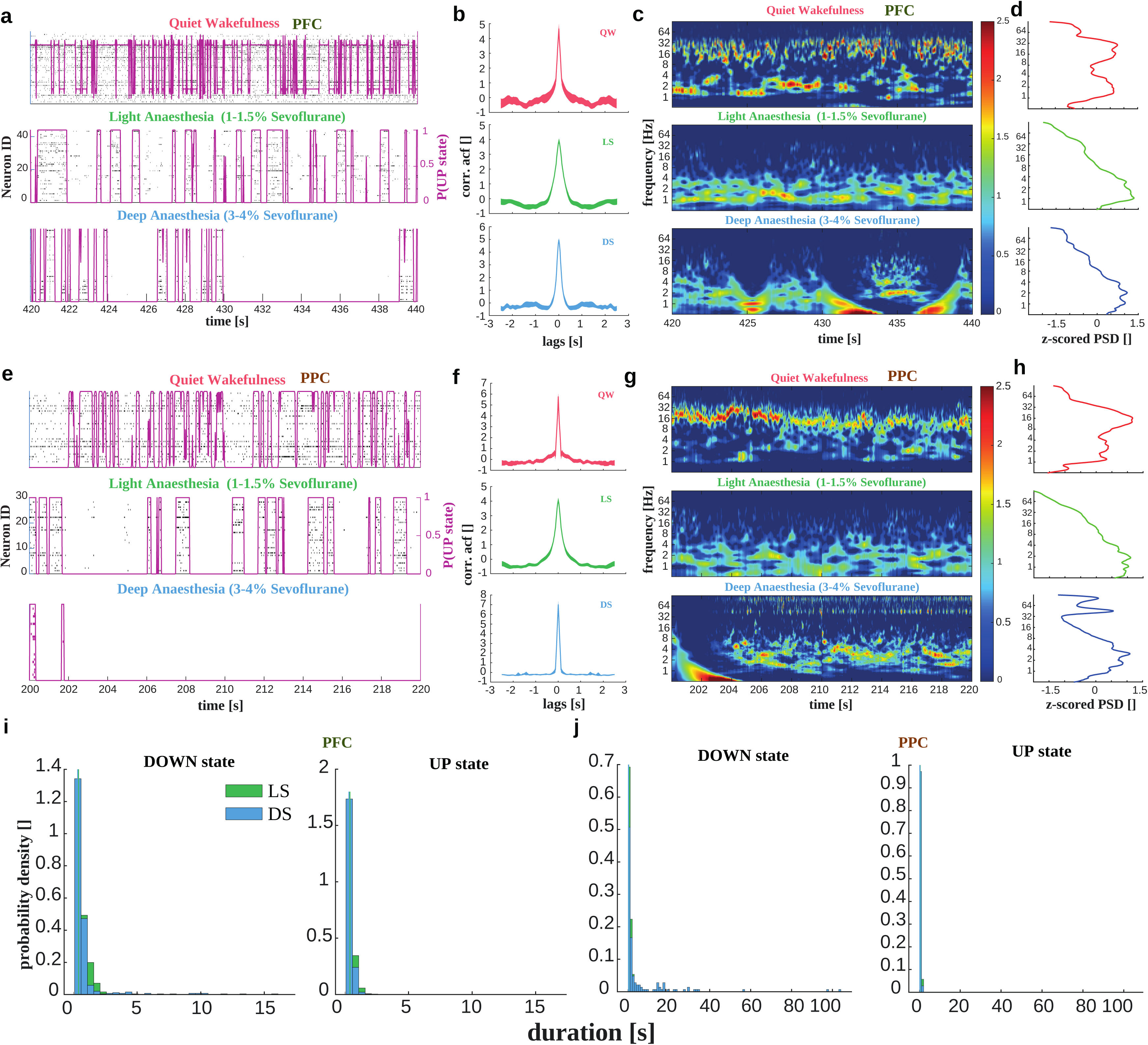
PFC and PPC display distinct neuronal population activity patterns under different conscious states. A. PFC - Reorganisation of spiking activity into periods of UP and DOWN states in anaesthesia (LS - Green, DS - Blue) as compared to desynchronised activity during QW (Red). DS also displays extended periods of cortical silence (seconds to minutes) termed “burst suppression”. B. PFC - Autocorrelograms of the population firing vector computed in chunks. While LS shows a wider peak, the strength of the non-shifted correlation is highest during DS and lowest during QW. C. PFC - z-scored time-frequency wavelet decomposition of the corresponding LFPs. While QW is identified by its characteristic ongoing high-frequency power which follows spiking, this is suppressed during both types of anaesthesia, where slower rhythms dominate. The purple line indicates the learned UP state path from the 2-state HMM. *n.b.* - Shown is one example period from Monkey A. D. PFC - z-scored spectrum for the example epochs. While all frequencies are well-represented during QW, especially beta and gamma, during LS and DS, the spectral activity is concentrated in the 0.1-8 Hz range. E, F G and H depict the same as above, but for the PPC. I. Col 1 & Col 2 - Duration of DOWN and UP states in the PFC. Col 3 & Col 4 - Duration of DOWN and UP states in the PPC. Green is LS, Blue is DS. While during both LS and DS, the durations of the up states stay below 3s, the durations of the silent periods extend to 15s (PFC) and 100+s (PPC) during DS.

We quantified the observed pattern of activity during these resting state periods (in both wakefulness and two levels of anaesthesia) using a two-state Hidden Markov Model (HMM) fit to the population firing vector. From the learned state sequence, the durations of bursting (UP) and silent (DOWN) periods were computed. While the model did not fit wakefulness well (Fig. 2A and 2E top row - purple line P(UP)), indicating the absence of such states during this period, it detected UP/DOWN states reliably in both LS and DS, in both PFC and PPC (Fig 2A and 2E, middle (LS) and bottom (DS) rows). While in Monkey A (PFC) and Monkey C (PPC), DS was typically characterised by long silent periods up to ∼16s (PFC, Fig. 2I Column 1) and ∼100s (PFC, Fig. 2I column 3), silent periods in both LS and DS were restricted to ∼2.6s in Monkey B (PFC, SIFig. 3)

**Figure 3.**
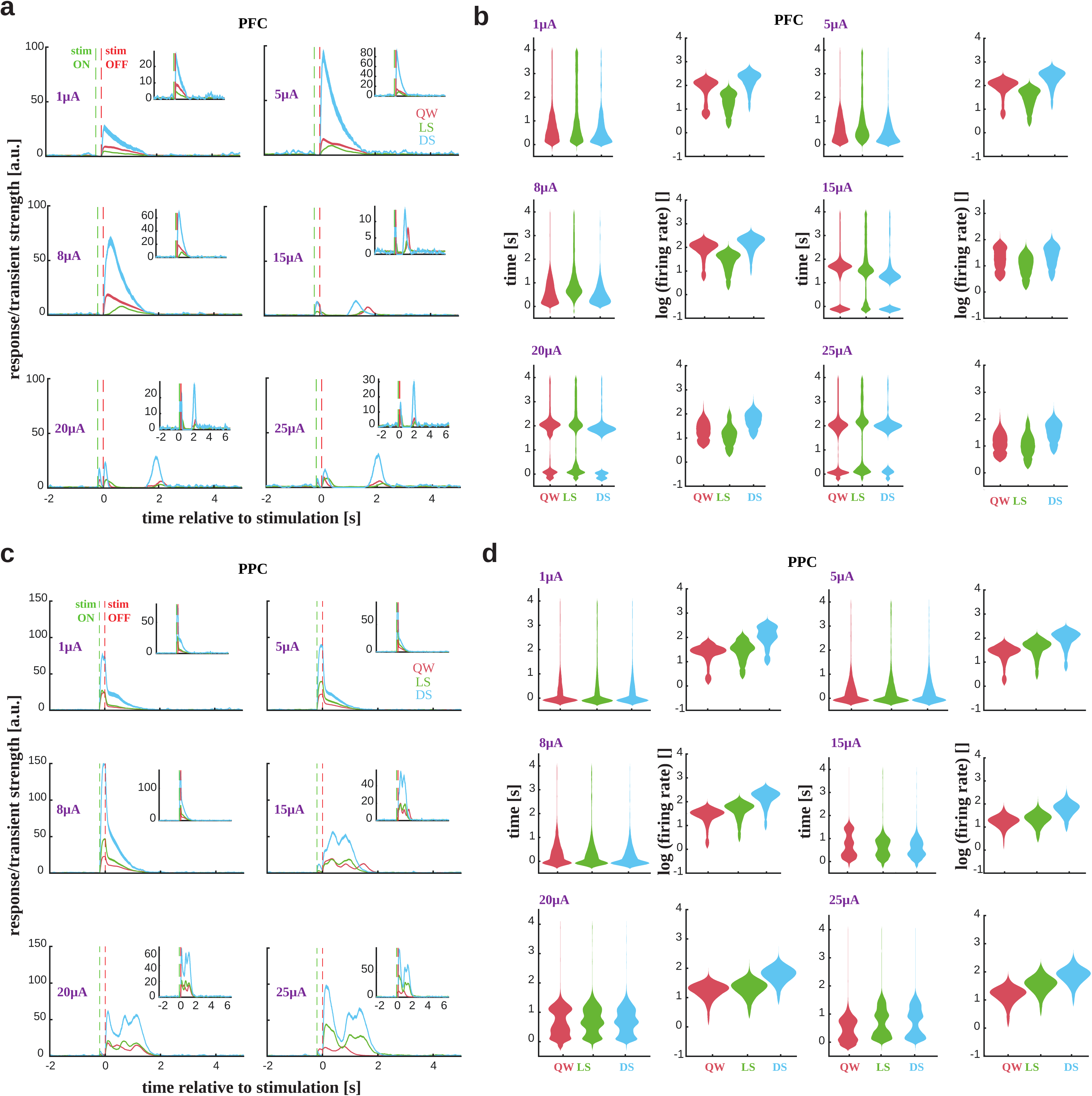
Spiking characteristics. A. Peri-stimulus time histograms (PSTHs) computed for all stimulation amplitudes (1, 5, 8, 15, 20, 25µA) during QW, LS and DS, in the PFC. The firing rates were averaged over channels and trials, and aligned to stimulation offset. Insets are zoomed in on the response shape. B. This panel characterises the above activity as violin distributions, estimated using an n-peak detection algorithm. It represents the times at which the peak transients and rebound occurred. While the cortical population response to low amplitude stimulation is temporal fast, transient, and exhibits an initial inhibition followed by excitation, higher amplitudes elicit an initial transient followed by long lasting inhibition, after which the discharge activity rebounds. C. Same as A, for PPC. D. While the cortical population response to low amplitude stimulation is temporal fast, transient, and exhibits an initial inhibition followed by excitation, higher amplitudes elicit an initial transient followed by short inhibition, after which the discharge activity rebounds, followed by another rebound. This does not occur in the PFC.

Finally, a synchrony index (SI) computed from the positive half of the autocorrelation function (ACF) excluding the peak suggests that while synchrony of population firing is relatively high during both LS and DS periods, QW displays low synchrony (Fig. 2B (PFC) and 2C (PPC); SIFig. 1), in both the PFC and the PPC. The difference in synchrony between each state was evaluated statistically, and all differences were significant with all p_Comp_ < 0.001. We attribute this to the coherent UP/DOWN states observed under anaesthesia.

### State dependence of cortical responsivity to electrical microstimulation

Following the characterisation of prefrontal and parietal states during wakefulness and different depths of sevoflurane anaesthesia, we examined the pattern of cortical response to electrical microstimulation administered at the centre of the implanted Utah array during these different states. Trial and channel averaged peri-stimulus time histograms (PSTHs) for spiking activity (Figs. 3A - PFC & 3C - PPC), and the same for LFPs (0.1 to 250Hz) (Fig. 4A - PFC & 4C - PPC) were computed aligned at stimulation offset.

**Figure 4.**
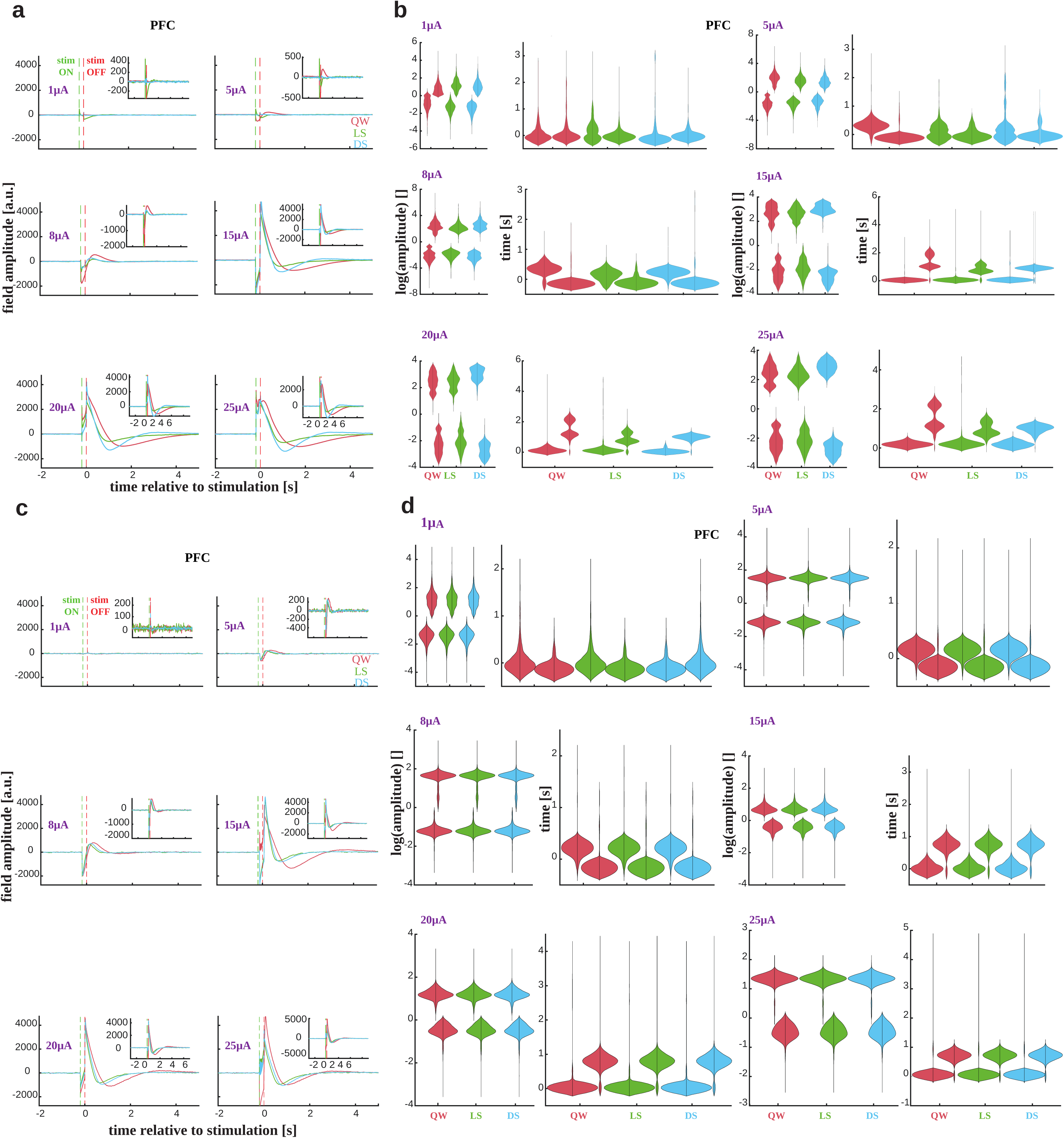
LFP characteristics. A. LFP activity for all stimulation amplitudes (1, 5, 8, 15, 20, 25µA) during QW, LS and DS, in the PFC. Insets are zoomed in on the response shape. B. This panel characterises the above activity in terms of the timing of the induced up or down state post stimulation. While the cortical population response to low amplitude stimulation first elicits a negative deflection followed by a positive overshoot, this pattern is reversed for higher amplitudes. C. Same as A for PPC D. Same as B for PPC (*n.b.* - In both the LFPs and neuronal discharges, the heterogeneous response patterns can be grouped into two distinct groups, viz. those in response to “low” amplitudes (1,5,8 µA) and those in response to “high” amplitudes (15, 20, 25 µA).)

For lower microstimulation amplitudes viz. 1, 5 and 8 µA, the baseline-normalised spiking activity displayed a suppression during the microstimulation period, followed by a sharp excitatory transient immediately after the end of microstimulation for all consciousness states (QW, LS and DS), in both PFC and PPC. Perhaps counterintuitively, the excitatory transient was stronger in amplitude for DS compared to QW and LS, in both recorded regions. However, for higher amplitudes (15-25 µA), we observed a different pattern present again in all consciousness states. In the PFC, spiking activity increased during the microstimulation period, which was then followed by a period of long-lasting inhibition (∼1-1.5seconds), finally bookended by a later rebound. This rebound was the lowest during light anaesthesia, and highest during deep anaesthesia in the PFC (Fig. 3A). Interestingly, while only one rebound occurred in the PFC, two rebounds manifested in the PPC, with the inhibition and rebound occurring faster and lasting for a shorter time (Fig. 3C). Furthermore, both the transient and the rebounds in the PPC followed a gradient according to the state, i.e. DS > LS > QW.

The above-mentioned characteristics are summarised as violin distributions (Fig. 3B - PFC, and Fig. 3D - PPC), estimated using an n-peak detection algorithm. In the PFC, for the lower stimulation amplitudes, the response during QW was the fastest, followed by during DS, with the slowest response manifesting during LS (Fig. 3B). However, in the PPC, the response occurred earlier in LS than during QW, with DS being the fastest. (Fig. 3D). Furthermore, the strength of the responses was highest during DS, which can potentially be attributed to increased cortical membrane sensitivity during burst suppression periods^80,81^. While in the PFC, the response and rebound strengths were weakest during LS, with DS being the highest, in the PPC, again, the response strength showed an increasing trend with anaesthesia. The mean occurrence times, their strengths, and their dispersal, under QW, LS and DS, for both Monkey A (PFC) and Monkey C (PPC) are catalogued in SI Table 1 (PFC) and SI Table 2 (PPC), along with confidence intervals (CIs) for the difference in means (If a CI includes 0, then no significant differences between the two states are seen).

The LFPs also displayed stimulation-dependent heterogeneity in the response pattern and strength, both in the PFC (Fig. 4A) and the PPC (Fig. 4C). For lower amplitudes (1, 5 and 8 µA), following stimulation offset, a negative deflection is seen, often followed by a positive rebound. However, these responses are noisy (Fig. 4C insets), and highly variable. For higher amplitudes (15-25 µA), following stimulation offset, a strong positive deflection is elicited followed by a delayed negative dip. These responses are highly stereotypical and consistent (Fig. 4C insets). While the first transient happened earliest during QW, and latest during DS, the reversal occurred latest during QW and earliest during DS, in both the PFC and the PPC. Furthermore, the initial response transient was sustained for the longest time (peak width duration) during QW, while it resolved itself the fastest during LS, pointing to its role in modulating membrane potentials of the constituent neurons of the population in the slow rhythm range (∼0.5-2Hz; data not shown). The mean occurrence times, their strengths, and their durations, along with their dispersal, under QW, LS and DS, for both Monkey A (PFC) and Monkey C (PPC) are catalogued in SI Table 3 (PFC) and SI Table 4 (PPC) respectively, along with confidence intervals (CIs) for the difference in means (If a CI includes 0, then no significant differences between the two states are seen).

In summary, these results show that causal perturbation via targeted electrical stimulation elicits state, stimulation strength, and space dependent heterogeneity in the response activity of the fronto-parietal network.

### Spatiotemporal modulation of field activity in different consciousness states

To study whether the state of consciousness has an effect on lateral signal propagation in the two cortical regions, the electrodes on the Utah array were binned according to their distance from the stimulated electrodes in 0.4mm bins, from the nearest (0.4 to 0.8mm) to the farthest (2-2.4mm in PFC and 2.4-2.8mm in PPC). Responses to ICMS were pooled for all stimulation amplitudes in Monkey A and Monkey C, while in Monkey B, only the response to 15 µA stimulation was analysed (only 2 amplitudes were tested in Monkey B viz.15 and 100 µA. 100uA displayed complete suppression in Monkey B, and thus was discarded from the analysis). The field response profiles by distance to the stimulated electrode are displayed in Fig. 5A (PFC) and 6A (PPC) respectively, with 2 stimulation amplitudes being represented, viz. 8 and 15 µA.

**Figure 5.**
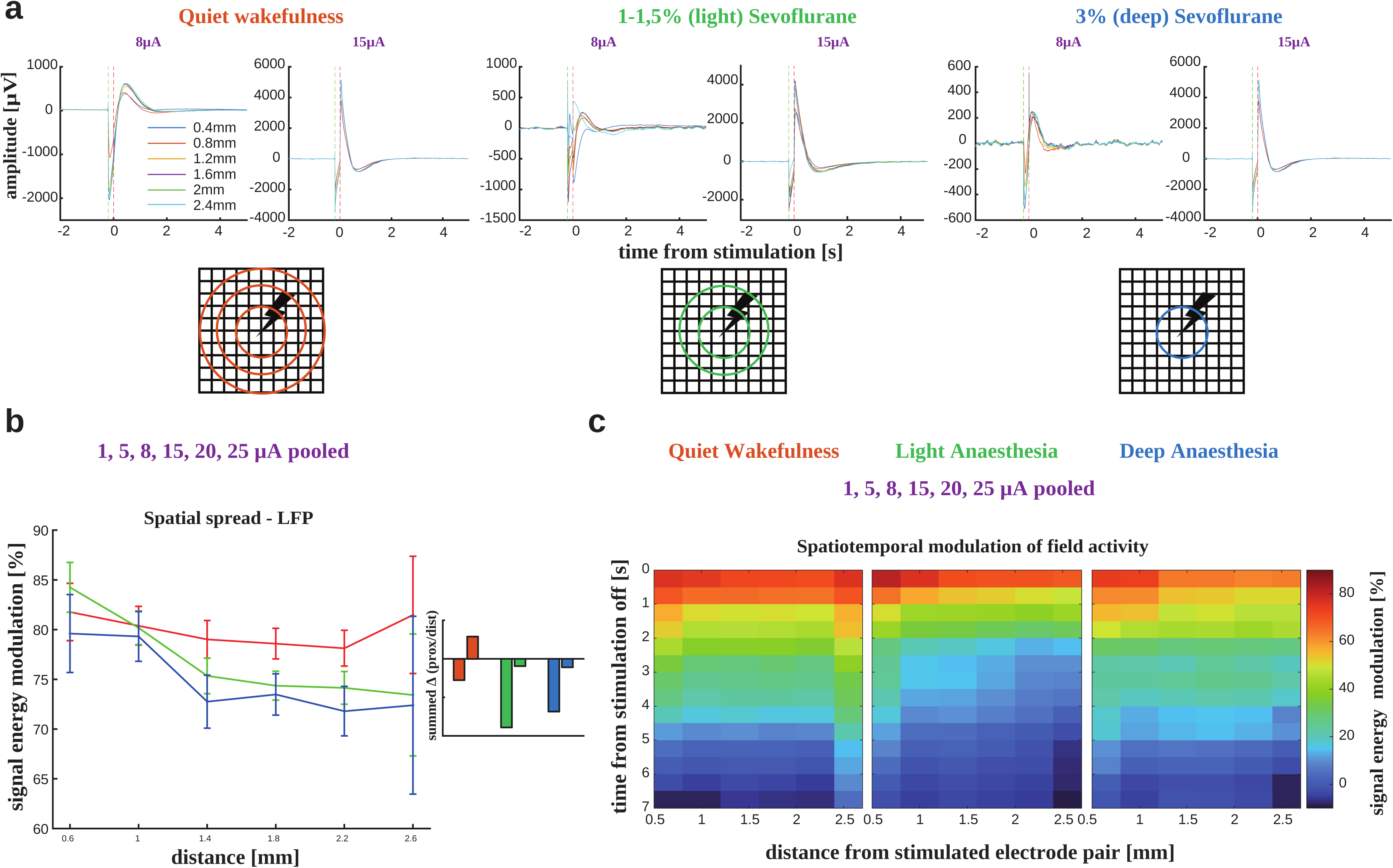
Modulation of cortical information flow in different states - PFC. A. Shows the average LFP response aligned to stimulation offset in the three states, viz. quiet wakefulness, light and deep anaesthesia, for two example amplitudes from each regime, i.e. “low” and “high” stimulation, sorted by distance, from nearest (0.4mm) to farthest (2.6mm) nodes from the centre. As noted previously, stimulation was delivered to the two centre electrodes. B. Whereas during QW, proximal and distal populations show similar modulation of field activity, this modulation falls off over space during both types of anaesthesia, with the decrease being stronger during DS (Insets show the sum of the first derivative for the proximal and distal populations). Data points show the mean and SEM. C. Top - Shows the evolution of field signal energy modulation in 0.5s bins starting immediately after stimulation offset, until 7s after stimulation. While the non-monotonicity is preserved in almost all time bins during QW, it does not manifest during LS or DS at all, showing a lack of spatial spread of information.

Because of the distinct response patterns to the low and high amplitude regimes elicited in the spiking activity, viz. a simple transient excitation versus a transient followed by inhibition, and rebound, in both the PFC and PPC, we restricted the analysis of the spatial modulation of population activity to the LFPs.

LFP signal energy modulation over distance was computed as the normalised pre/post change in activity in a 4s bin following the end of stimulation. The signal response was modulated strongest during QW, displaying non-monotonicity over distance, while they were modulated the least during DS, and decreased sharply over distance (Fig. 5B). Neuronal activity in LS displayed modulation strength in between QW and DS, and also decayed linearly over distance, albeit slower than DS (Fig. 5B). This points to a maintenance of the strength of modulation at longer distances during QW, with proximal and distal populations being modulated equally, while signal propagation is restricted with increase in distance, in both states of anaesthesia. These are confirmed by the analysis of the sum of the rate of change over distance, combined with a Bayesian bootstrapped comparison-of-means. The spatiotemporal characterisation for Monkey B is presented in SI Fig. 4.

In the PPC however, modulation decreased with distance in all three states pointing to no non-monotonic recovery of signal propagation with distance (Fig. 6B). However, as opposed to the PFC where in the LFPs, the strongest modulation was observed during QW (Fig. 5B), in the PPC, the strongest modulation was seen in the DS.

**Figure 6.**
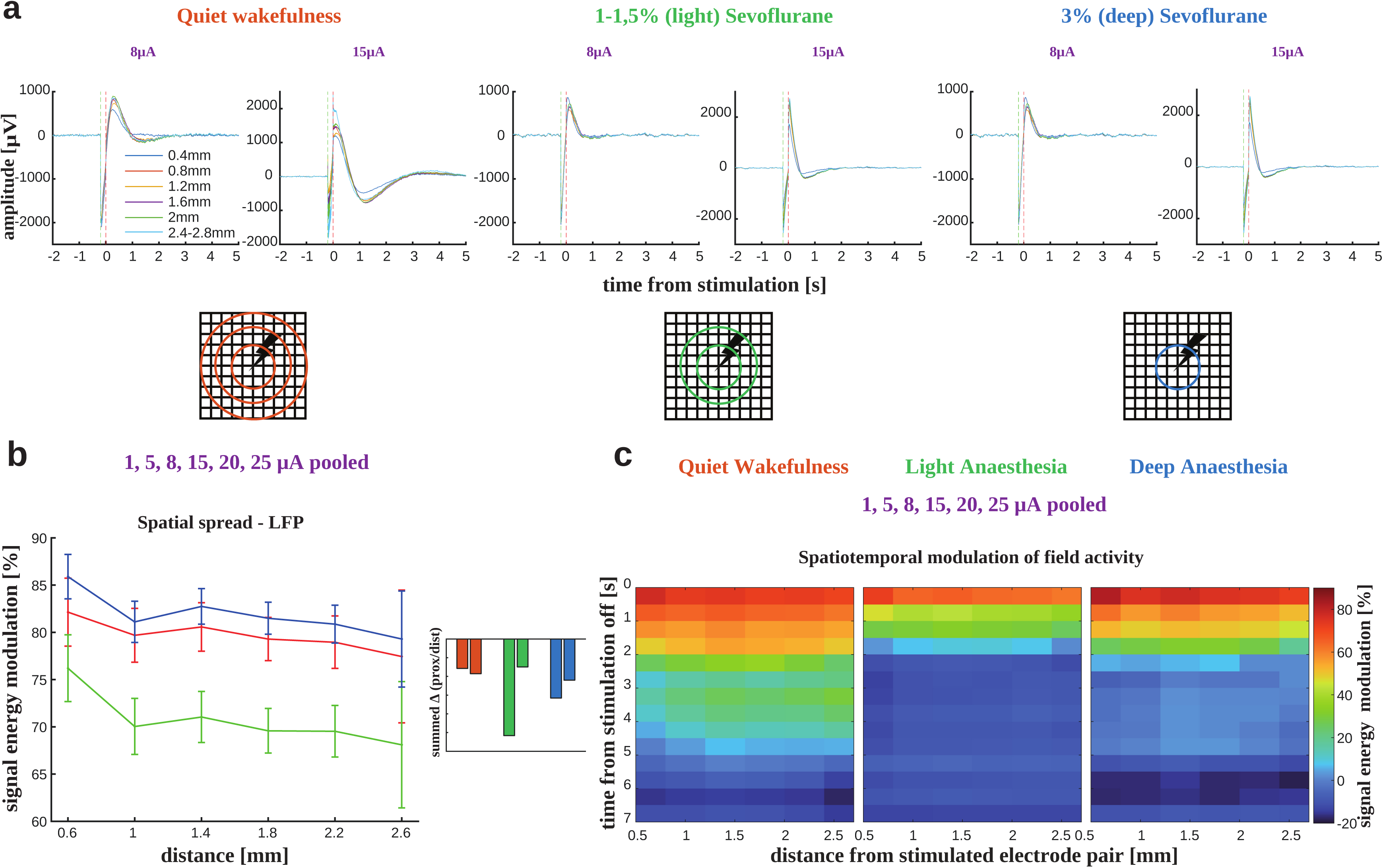
Modulation of cortical information flow in different states - PPC. A. Shows the average LFP response aligned to stimulation offset in the three states, viz. quiet wakefulness, light and deep anaesthesia, for two example amplitudes from each regime, i.e. “low” and “high” stimulation, sorted by distance, from nearest (0.4mm) to farthest (2.8mm) nodes from the centre. As noted previously, stimulation was delivered to the two centre electrodes. B. Field signal energy modulation in the PPC shows that no non-monotonic structure is observed at all in the LFPs in the PPC, as opposed to the PPC. Furthermore, As opposed to the PFC, in the PPC, DS modulation remains higher than QW and LS. C. While the strength of modulation persists for longer after stimulation in the PPC during QW, it extinguishes fastest during LS, and then DS. No non-monotonic structure is observed in any time-bin in any state after stimulation.

A 0.5s bin by bin analysis of this modulation in time in the PFC, starting immediately after stimulation offset, displays the preservation of this non-monotonicity during wakefulness (Fig. 5C; Left Column), even up to 6s after stimulation offset, in the LFPs. However, during both levels of anaesthesia, the drop off of signal energy modulation over distance remained in all time bins (Fig. 5C Middle and Left Columns). In the PPC, the decay in time was even faster, strong excitatory modulation extinguishing already after ∼2.5s (Fig. 6C).

Taken together, our results suggest that the prefrontal cortex not only displays distinct differences in responses in spiking activity and LFPs to electrical microstimulation, it also displays differential responsivity to low and high amplitudes of stimulation, under different states of (un)consciousness. Most importantly, our data show that while field signal propagation in the PFC is starkly reduced under anaesthesia, as compared to wakefulness, only the strength of modulation displays significant differences between the states in the PPC.

## Discussion

The PFC exhibits a rich diversification of structure^82–84^, and consequently, function. It serves as an integrative hub for a multitude of cognitive processes such as inhibitory control, decision making, cognitive control, working memory, and the integration of sensory information with planned actions^85–91^. Under anaesthesia, both rodent and human studies have revealed notable alterations in PFC activity. Specifically, anaesthesia has been shown to induce a transition from desynchronised to highly synchronised oscillatory activity in the PFC of rodents^72,92,93^. In humans, anaesthetic agents have been observed to disrupt functional connectivity within the PFC, which correlates with a decline in conscious awareness and cognitive function^43,94,95^. Pressingly, what can we learn from characterising macaque PFC population dynamics by monitoring and perturbing neuronal populations during these (un)conscious states?

### Altered mesoscale cortical dynamics under anaesthesia

Previous studies have monitored cortical populations across the dorsal and ventral pathways in different animal models during rest. Furthermore, global dynamics have been studied using gross-scale techniques in humans. Perturbational studies in humans and monkeys under different (un)conscious states have been performed with both DBS and TMS^73,96–99^. However, to date, consistent, simultaneously-recorded mesoscale cortical populations in different states have not been monitored, and perturbed.

We show that chaotic and dispersed spiking activity in the same population during the awake state reorganises into coherent UP/DOWN states during anaesthesia, both during light and deep anaesthetic states (Fig. 2). The distinguishing feature between LS and DS both in the PFC and PPC seems to be a long tail of the duration distributions of the down states during DS (Fig. 2), thus signifying a quieter, i.e. less active cortex. However, a higher population synchrony in DS agrees well with the significantly larger low-frequency power (Fig. 2, delta and theta), and therefore coherence (data not shown). These long periods of cortical silence, termed burst suppression in neurological EEG literature, could reset cortical responsivity during anaesthesia at the mesoscale.

### Response characteristics of the PFC and PPC

In the past, a majority of causal perturbational studies have been used either to bias perception and behaviour by stimulating stimulus-selective neurons in sensory cortices^74,100–104^ or object recognition areas^105^, or for brain computer interfaces (BCIs)^106^, and therapeutic purposes^107–109^. While studies that electrically and pharmacologically perturb the PFC in humans^36,110^ and animal models^10,111^ exist, they are sparse and not systematised, neither in the behaviour or sub-behavioural range, nor under different arousal states. Our study addresses this *lacuna* specifically with respect to intracortical microstimulation by assaying the primate higher associational cortices. Crucially, no studies exist that simultaneously monitor and perturb the same consistent population of prefrontal and parietal neurons under different states of consciousness, thus targeting the fronto-parietal loop.

Our results (Figs. 3A and 3B) suggest the existence of two membrane-responsiveness regimes with respect to electrical challenge in these cortical populations. We show that low amplitude stimulation seems to display an initial inhibition followed by excitation (Fig. 4A - Left). However, large current injections elicit first a strong transient, followed by a long-lasting inhibition, after which the spiking activity rebounds with a vengeance (Fig. 4A - Right). Inhibitory neuronal populations, though comprising only 15-30% of the cortex^112,113^, tightly regulate microcircuit function^114,115^. Post-inhibitory rebound and its relationship to recurrent inhibition is widely-known in a variety of intrinsically and extrinsically-driven contexts^116^. Large current insults could reflect a disruption of this inhibitory control, and its subsequent release, pointing to an inhibitory population led network reorganisation^117,118^. This heterogeneity, also preserved in the LFPs (Figs. 4A and 4B), could therefore point to downstream perisynaptic modulation, allowing the cortical membrane potential to display differential activity as a consequence.

In the PPC, in addition to the transient-inhibition-rebound, we observe a second rebound that happens very close to the first (Fig. 3C). Because the firing rate before this rebound does not return to baseline, and instead manifests as a “second peak”, this extra rebound could potentially arise from recurrent excitation, and could be crucial in self-generated and sustained neural activity, that can be broadcasted to organise on-line circuit control^119,120^.

### The responsivity paradox

Our data also indicate that fronto-parietal responsivity is highest during deep anaesthesia. What seems at first paradoxical, lends itself to a remarkable interpretation in terms of equilibrium dynamics. During deep anaesthesia, long periods of iso-electric voltage lines (in the EEG) interspersed with brief, large fluctuations exist (Fig. 2A), which is termed “burst suppression”^121^. We speculate that during these extended periods of cortical silence, the neuronal membrane is hyper-sensitive^80^, thereby allowing it to return to equilibrium, during which, any reasonably strong perturbation could potentially push the system into an accelerated, hyperbolic trajectory, manifesting as fast and strong responses. Such empirically-determined cortical activity patterns under perturbation, rest and different states are imperative for constraining models of brain function at multiple scales.

Contrasting resting state activity profiles with the observed response dynamics underscores the potential limitations in inferring the exact state of consciousness (for e.g., depth of anaesthesia) from temporal patterns in population activity. Despite the varying distributions of quiescent intervals across sessions and subjects, the pattern of increased responsivity coupled with limited spatial propagation during deep sedation (DS) was a consistent finding. This observation is bolstered by the analysis of proximity to criticality (see SI. Fig 1), which showed uniform spatial and temporal response characteristics across sessions and subjects, regardless of any significant shifts in criticality between light sedation (LS) and DS.

Moreover, the hypothesis that direct stimulation may induce prolonged suppression, culminating in a synchronised and markedly amplified rebound, suggests that during DS, complexity indices - evaluated through methods like perturbational complexity index (PCI) and pattern analysis - would be reduced due to this enforced, homogenous, and intensified rebound effect. Consequently, beyond a certain depth of anaesthesia, where neuronal populations reach a threshold of inhibitory saturation, a progressive decline in criticality or complexity metrics may not be evident. In effect, these metrics potentially only reflect this specific inhibitory population led dynamics, rather than a true reconfiguration of information, and thus complexity. Future investigations could explore these dynamics through BOLD-fMRI under varied anaesthetic conditions and DBS protocols, coupled with advanced PCI evaluations using complex TMS protocols, including pulse-train stimulation, to thoroughly examine and correlate activities across different scales.

### Signal communication is maintained during wakefulness in PFC but not in PPC

Anaesthesia is speculated to induce unconsciousness in many ways^47,60,122^, a primary one among which is by suppressing information processing and communication. This is evidenced by global inhibition, i.e. the reduction in neuronal spiking activity^123,124^. Additionally, local synchrony increases but global synchrony has been observed to be disrupted under anaesthesia^125^, while during wakefulness, although locally neurons remain uncorrelated in their firing, large-scale synchrony is observed across brain regions^126–128^, specifically with the persistence of the Default Mode Network (DMN)^129^. This change in synchrony patterns under anaesthesia suggests a disruption in normal information processing in the brain, which at the prefrontal cortical level is confirmed by our results.

Crucially, the dorsal and ventral PFC have been shown to contain features that distinguish them from lower cortical regions^130^. For example, instead of distinct organisation into cortical columns, the PFC instead shows greater lateral connectivity among distributed pools of functionally distinct neuronal cell types^83,131–133^. Layer 5 pyramidal neurons project out of the cortex and receive extra-cortical and subcortical efferents to establish overarching connectivity^134,135.136^. Long-range fronto-parietal connections exist which form the fronto-parietal loop, a unit central to the GNW^14,16,137^. Most importantly, the PFC displays not only abstract tuning such as task-phase^138^, but also stimulus features^139^. These various modes of differentiation and integration^70^ are ideal for complex information processing, such as those required for conscious awareness^13,140^. A unique and computationally significant property of neuronal networks is functional connectivity, which can be measured using noise correlations at the mesoscale^141,142^. The structure of this functional connectivity in lower cortical areas is a linear decrease with lateral distance, both under wakefulness and anaesthesia, whereas in the PFC it is non-monotonic, showing that proximal and distal populations are equally connected^143,144^. This is also evidenced by patchy anatomical connectivity between pools of excitatory and inhibitory populations^83,131,133,145^. This unique pattern of connectivity in the PFC suggests that it can be used for complex integrative functions^87,146–148^ that lower order areas cannot perform. Interestingly, primary sensory areas, with the exception of the somatosensory cortex, aren’t much perturbed during anaesthesia, even at deep levels^149,150^. These observations lend support to the hypothesis that loss of consciousness is triggered by a lack of higher-order integration in the cortex^66,151^. Lower areas don’t need long-range functional connectivity because they process low-level stimulus features independently and don’t need to integrate. However, feature-binding (Gestalt) is necessary for conscious perception^152–156^, the PFC being a prime candidate^157–159^.

From the standpoint of GNW, although the PPC is indeed part of the fronto-parietal loop^43^, and distinct patterns of directionality, strength and functional organisation are observed across states of consciousness in each individual module of the loop^42,43^, it is the PFC that is supposed to amplify, broadcast and coordinate brain-wide activity to allow for consciousness to emerge 15^15,17^. Therefore, we posit that anaesthesia must necessarily disturb the PFC, not that it is only sufficient to dampen PFC or PPC activity. If anaesthesia causes an information bottle-neck, long-range coordination of neuronal activity by an orchestrating node would become weak or impossible. Our results (Fig. 5) show – that although fast spatial spread of global activity in V1 is seen immediately post ignition – that strong horizontal connectivity and lateral information flow, only in the PFC, is a signature of conscious wakefulness and indeed breaks down under anaesthetic unconsciousness, suggesting that such complex functional cytoarchitecture in frontal regions is necessary for integrative processes crucial to conscious awareness, which subsequently implicates its disruption in loss of consciousness.

### Causal perturbation of the fronto-parietal circuit and its systemic implications

In the ongoing discussion surrounding the role of the PFC in consciousness and DoCs, studies employing intracranial electrical stimulation (iES) to causally perturb networks of interest have recently begun to not only provide unique, unexpected and contradictory insights^110^, but also elicit spirited debate^36^. Most importantly, studying cortical response properties under external stimulation can help elucidate the mechanistic underpinnings of consciousness, as opposed to purely correlative studies. Causal perturbational studies are crucial as they allow for the manipulation of specific neural populations, thereby providing direct evidence of their role in consciousness. To even begin to design appropriate experiments addressing this large gamut of questions, it is vital that a foundation is securely built on thorough electrical and optical characterisation of these mesoscale networks. Furthermore, insights gained from such studies can be used to constrain larger neuronal models of cortical activity under different states of consciousness.

Studies employing iES or ICMS in animal models have also provided indirect support for a central role of the PFC in conscious awareness. For example, it has been shown that when the primary visual cortex (V1) is stimulated, correct detection trials consistently display sustained PFC activity as opposed to misses^75,76^. In terms of the state of consciousness, a landmark study in rodents demonstrated robust RoC after carbachol injection to the PFC, but crucially, not the PPC^10^. Interestingly however, when the thalamus is stimulated using deep brain stimulation (DBS) to arouse the experimental subject, both the PFC and the PPC are strongly activated, indicating the engagement of fronto-parietal populations during RoC^44,160,161^. In task contexts for instance, global deviants in the local-global task^162^, which are a signature of conscious access, can then be decoded from the PFC when other behavioural signs of arousal re-manifest^44^. DBS has also shown promise in “enhancing” conscious access in patients with minimally conscious state (MCS), marking a significant advancement in DoC intervention^96^. These findings trace a functional link between the thalamic relay and higher associational populations in conscious access, potentially engaging a broader thalamo-cortical circuit, thereby restoring consciousness in patients with DOCs, while also providing a more nuanced approach in identifying potential therapeutic targets, such as cautiously employing direct cortical stimulation - rather than strongly invasive and high-strength DBS - could. However, systematic protocols for characterising ICMS effects during wakefulness or anaesthesia (as a proxy for DoCs) are lacking. Our study attempts to fill this gap by exploring myriad stimulation protocols and parameters (Fig. 1).

### Limitations

In our study, we recorded stable prefrontal and parietal populations during all states of (un)consciousness. However, it is prudent to note that the PFC and PPC data come from two different animals, due to the unfortunate loss of the PFC and the PPC array in each animal respectively (COVID-19 *etc*; in the other animal, no spiking activity could be recorded from the PPC array).

This inherently limits the direct comparison of the two areas forming the frontoparietal loop. However, it is our firm belief that the consistency of responses and functional modulation over space and time seen in our data would predict the same across simultaneous recordings too.

Furthermore, since we used stimulation regimes that have previously been shown to elicit behavioural responses^75,76,163–165^, we did not probe the cortex classically with single-pulse stimulation, as our goal was to also investigate response characteristics and signal spread under these un(conscious) states.

## Conclusion

Taken together, our results reveal that the PFC and the PPC exhibit unique and intriguing responses when subjected to electrical stimulation, specifically shedding light on the putative functional role of the PFC in regulating states of consciousness. Most importantly, we show the restriction of lateral signal propagation under anaesthesia in the PFC, which could result in break-down of crucial integrative processes that are necessary for consciousness and cognition. These results can further significantly inform and constrain cortical models of brain function in health and disease.

Further investigation into the effects of elevated electrical stimulation intensities on these higher order association areas offers a promising avenue for future research. By delving into the consequences of stimulation whose parameters are in the range to elicit behavioural effects, we have the opportunity to elucidate the nuanced relationship between PFC activity and specific behavioural metrics, particularly in arousal and its loss. This line of enquiry holds substantial clinical implications, especially for individuals afflicted with DoCs. Given that our current stimulation design did not demonstrate behavioural signs of arousal (this is demonstrated with a different stimulation regime, in forthcoming work) or long-range cortical excitability, we hypothesise that higher stimulation intensities or alternative stimulation patterns, as guided by dynamic brain activity^100,166,167^, could potentially increase cortical excitability and thus awaken subjects under anaesthesia. To this end, we are initiating further studies aimed at assessing the feasibility of reversing anaesthesia-induced unconsciousness through targeted electrical stimulation of the frontoparietal loop.

## Acknowledgments

We thank Dr. Lynn Uhrig for her assistance with anaesthesia. We are thankful to Jeremy Bernard for assistance with the custom-made setup for the experiment. We would also like to thank Prof. Dr. B S Dwarakanath for helpful discussions and comments, and proof-reading assistance.

## Author contributions

Conceptualisation: AD, TIP; Data curation: AD; Formal analysis: AD; Funding acquisition: BJ, SD, TIP; Investigation: AD, MKA, MG, MR, RHV, BJ, TIP; Methodology: AD, MKA, RHV, TIP; Project administration: TIP; Resources: MG, MR, BJ, SD, TIP (lead); Software: AD; Supervision: BJ, TIP (lead); Visualisation: AD (lead), RHV (supporting); Writing - original draft: AD; Writing - review & editing: AD, MKA, MG, MR, RHV, BJ, TIP (lead).

## Funding

This work received funding from Inserm (BJ), CEA (TIP), Collège de France (SD), the European Union’s Horizon 2020 Framework Programme for Research and Innovation under the Specific Grant Agreement No. 945539 (Human Brain Project SGA3, T2.10; TIP), a grant from Templeton World Charity Foundation, Inc (TWCF0562) to TIP, and a grant from the Bettencourt-Schueller Foundation to BJ.

## Declaration of interests

The authors declare no competing interests.

## STAR Methods

### 1. Resource availability

Information and requests for data and specific code should be directed to and will be fulfilled by the lead contact or the first author – Theofanis I Panagiotaropoulos - (lead contact); Abhilash Dwarakanath -

### 2. Materials and availability

Not applicable

### 3. Data and code availability

Electrophysiological recordings used in this paper will be used in the future for other studies. Therefore, the data is available upon reasonable request; please contact the lead contact. Custom stimulation, data acquisition and extraction, preprocessing, and analysis scripts written in MATLAB are available on the first author’s GitHub - (https://github.com/abhilashdwarakanath/IntracorticalMicrostimulation, https://github.com/abhilashdwarakanath/-Un-ConsciousStates, https://github.com/abhilashdwarakanath/EphysPreprocessing).

The data used in this manuscript will be made available through the Human Brain Project’s EBRAINS Knowledge Graph, as a contribution from SGA3’s Task 2.10.

### 4. Experimental model and subject details

Two adult, male rhesus macaques (*Macaca mulatta)*, weighing between 10-12kg were recruited for the experiments. The subjects, denoted as Monkey “A” and Monkey “J”, were pair-housed in an enriched animal facility. They were trained to be chaired and head-posted in the recording chamber.

### 5. Methods details

#### Electrophysiological recordings

We performed extracellular electrophysiological recordings in the inferior convexity of the lateral PFC of 2 awake and anaesthetised monkeys using chronically implanted Utah microelectrode arrays^168^ (Blackrock Microsystems, Salt Lake City, Utah USA). We implanted the arrays 1 - 2 millimetres anterior to the bank of the arcuate sulcus and below the ventral bank of the principal sulcus, thus covering a large part of the inferior convexity in the ventrolateral PFC, which forms one node of the fronto-parietal loop. The arrays were 4×4mm wide, with a 10 by 10 electrode configuration and inter-electrode distance of 400μm. Electrodes were 1mm long therefore recording from the middle cortical layers. The monkeys were implanted with form-specific titanium head posts on the cranium after modelling the skull based on CT scans. All experiments were approved by the local authorities’ ethical committee (Protocol # - A18_028), and were in full compliance with the guidelines of the European Community (EUVD 86/609/EEC) for the care and use of laboratory animals.

#### Data acquisition and stimulation

Broadband neural signals (0.1–30 kHz) were recorded using Neural Signal Processors (NSPs) from Blackrock Microsystems Inc. Signals from the Utah array were digitised, amplified, and then routed to the NSPs for acquisition. The digitisation occurred on a CerePlex M Headstage which, coupled with Omnetics Y adapters, allowed us to simultaneously record and stimulate from the same electrodes. A CereStim 96 microstimulator from Blackrock Microsystems driven by MATLAB (2022a, MathWorks Ltd) code interfaced with their proprietary API was used to provide biphasic pulses between 1 to 210µA.

For the offline detection of action potentials, broadband data were filtered between 0.6 and 3 kHz using a second-order Butterworth filter (the filter was chosen such that it allowed a flat response in the passband while contributing the least phase distortion due to its low order, yet having an acceptable attenuation in the stop band, i.e. a roll-off starting at -20dB). The amplitude for spike detection was set to five times the median absolute deviation (MAD)^169^. Spikes were rejected if they occurred within 0.5 ms of each other, (multi-unit refractory period temporal threshold) or if they were larger than 50 times the MAD. Finally, all spikes that occurred at the same time point (within 3 samples at 30kHz) in a minimum of 60 channels (out of 96) were also discarded to exclude stimulation artefacts. All collected spikes were aligned to their minimum. For the analysis of LFP activity, the broadband signal was decimated to 500 Hz sampling rate using a Type I Chebyshev Filter, reliably preserving frequency components up to 200 Hz.

#### Experimental design

The animal was chaired and head-posted in the recording chamber, after which the eye-recording camera was aligned and centre-calibrated. The CereStim module was connected to the CerePlex M headstage for stimulation delivery, which was subsequently connected to the pre-amplifiers, which fed the recorded and referenced signals to the Neural Signal Processor (NSP) via fibre-optic cables.

The experiment consisted of four distinct phases viz. Quiet wakefulness (QW), induction and transition to anaesthesia, light anaesthesia (LS) and deep anaesthesia (DS). Each of the QW, LS and DS phases started with an initial recording of 15-20 mins of spontaneous activity. Next, multiple repeats (10 to 25) of a pulse train consisting of 50 pulses of amplitudes varying between 1 to 100µA were delivered to the two central electrodes, randomised, separated by 10-20s of dead-time. Each pulse was biphasic, being either anodal first, or cathodal first. Each phase lasted for 50µS with the inter-phase interval being 53µS. Therefore, each stimulation pulse lasted for ∼0.1995s. Following certain other stimulation protocols, a post-stimulation spontaneous phase of 15-20 mins was again recorded.

Transition to anaesthesia from wakefulness was achieved first by sedating the animal using a combination of Ketamine 1000 (3 mg/kg intramuscular) and Dexmedetomidine (15 µg/kg intramuscular). The animal was then released from head-fixation and carefully extracted from the experimental chair. The chair was then removed and a custom-built experimental bed was inserted and fixed. This bed was covered with a blanket and an inflatable heating pad was inserted. An injection of Antisedan (75 µg/kg intramuscular) was then given, as an antagonist of the sedative effect of Dexmedetomidine. In one experiment, the animal was laid in a prone position inside a stereotactic frame, while in the rest of the sessions, the animal was laid on its side. Next, the animal was intubated and an IV line was laid. Inhalation anaesthesia was initiated using 2% of Sevoflurane. The ventilator was switched on, and on confirmation of stable vitals, the door of the faraday cage was closed. After ∼80-90 minutes from the first injection, the LS experimental epoch was started by changing to and maintaining the anaesthesia at a low level of 1-1.5%. In one session, a paralytic agent, namely Curare, was used. After the end of the LS epoch, the inhalant anaesthesia was increased to 3% (DS). Crucially, we continued to record during this entire process thereby making sure that we were monitoring the same population of neurons throughout the 4 experimental phases.

At the end of the DS epoch, the inhalant anaesthesia was gradually reduced to 0. Upon awakening, the monkey was extubated, the IV line and rectal thermometer was removed, and disconnected from the ventilator and vitals monitor. Next, the animal was placed in the recovery box and returned to its cage. Finally, the animal was monitored for full recovery for the next 24 hours.

#### Evaluation of resting state activity

To statistically characterise the effects of the levels of anaesthesia, we fit a 2-state Hidden Markov Model (HMM) to the data. A firing rate threshold of 0.05 spikes/s (Monkey 1) or 2.5 spikes/s (Monkey 2) was set to exclude non-firing channels in the resting state periods, after which spikes were binned in 25ms (Monkey 2) or 50ms (Monkey 1) bins, and then summed and normalised to obtain the population firing vector. To this population vector for each state, k-means clustering was applied to obtain 2 distinct firing rate regimes. These cluster-centres were normalised and used as the initial emission probabilities (p(EM_init_)). Initial transition probabilities (p(TR_init_)) were drawn from a positive-normal distribution. An initial state sequence guess was obtained by setting all data above the mean to 2, and those below to 1. The HMM was trained on this data to obtain the learned p(EM_final_) and p(TR_final_). These learned probabilities were then used to generate the learned state sequence using the Viterbi algorithm, from which the durations of the UP and DOWN states were finally computed. This training and testing were performed using the HMM toolbox in MATLAB.

State time-frequency information was computed via a continuous wavelet transform using a generalised morse wavelet with 7 cycles and a time-bandwidth product of 3 x 30 at a sampling rate of 500Hz.

The measure of synchrony (Synchrony Index) was estimated as the sum of the positive-lag auto-correlation function (computed to a 0.5s lag) divided by the number of lags, after which it was subtracted from 1. We consider only the positive lags after 0, therefore we subtract the mean auto-correlation from the peak-value (i.e. 1 at lag-0 when the ACF is normalised).

#### Computing stimulation-related measures

To obtain the onset of each stimulation, the recorded pulse onset-offset analogue signal was differentiated, and zero crossings were detected. These onsets and offsets were sorted into their respective amplitude trials using the trial structure log file corresponding to each epoch. LFPs and spikes were collected for each trial and channel, and aligned to the stimulation offset. Resting state epochs were segmented from the recording using their respective epoch tags.

Mean responses to each stimulation amplitude were computed by concatenating the channel and trial LFPs and PSTHs and then averaging them. The errors were thus the standard deviation across these collapsed samples normalised by the square root of the number of channels, and finally multiplied by 1.96 to obtain the 95% CI.

Response characteristics of the neuronal discharge patterns, namely the first transient and subsequent rebound(s), and the LFP response, were characterised using MATLAB’s in-built *findpeaks* function. The dispersal was calculated as the distance to the cluster centre(s), which roughly corresponds to the variance of a normal distribution. For multi-peak rebound, k-means with n clusters was run to fit the underlying normal distributions.

LFP and spiking response characteristics comparisons were performed pairwise across state combinations (QW vs LS, QW vs DS, LS vs DS), using bootstrapped resampling. The differences were evaluated as the 95% confidence intervals (CI) on 10000 resampling runs.

### LFP modulation

where x(t) represents the LFP signal, k represents the trial index, and n represents the channel index.

The existence and strength of non-monotonicity in the spatial spread of field activity modulation was evaluated using a Bayesian bootstrap on 4 hypotheses, viz. flat, increasing, decreasing, and non-monotonic. Because the distances increase in even number pitch increments, the data and function were divided into “nearby” and “ far away” populations. The sum of the first differential was computed on these chunks. A 0 or positive index indicates non-monotonicity, whereas the magnitude of the negative sum denotes the steepness of the monotonic decrease. After 10000 bootstrapping runs, the hypothesis with the highest probability (above 0.25) was chosen as the most probable curve shape.

The MATLAB statistics and machine learning toolbox was used to implement all statistical analyses, with custom-written scripts.

---

**SI Figure 1.**
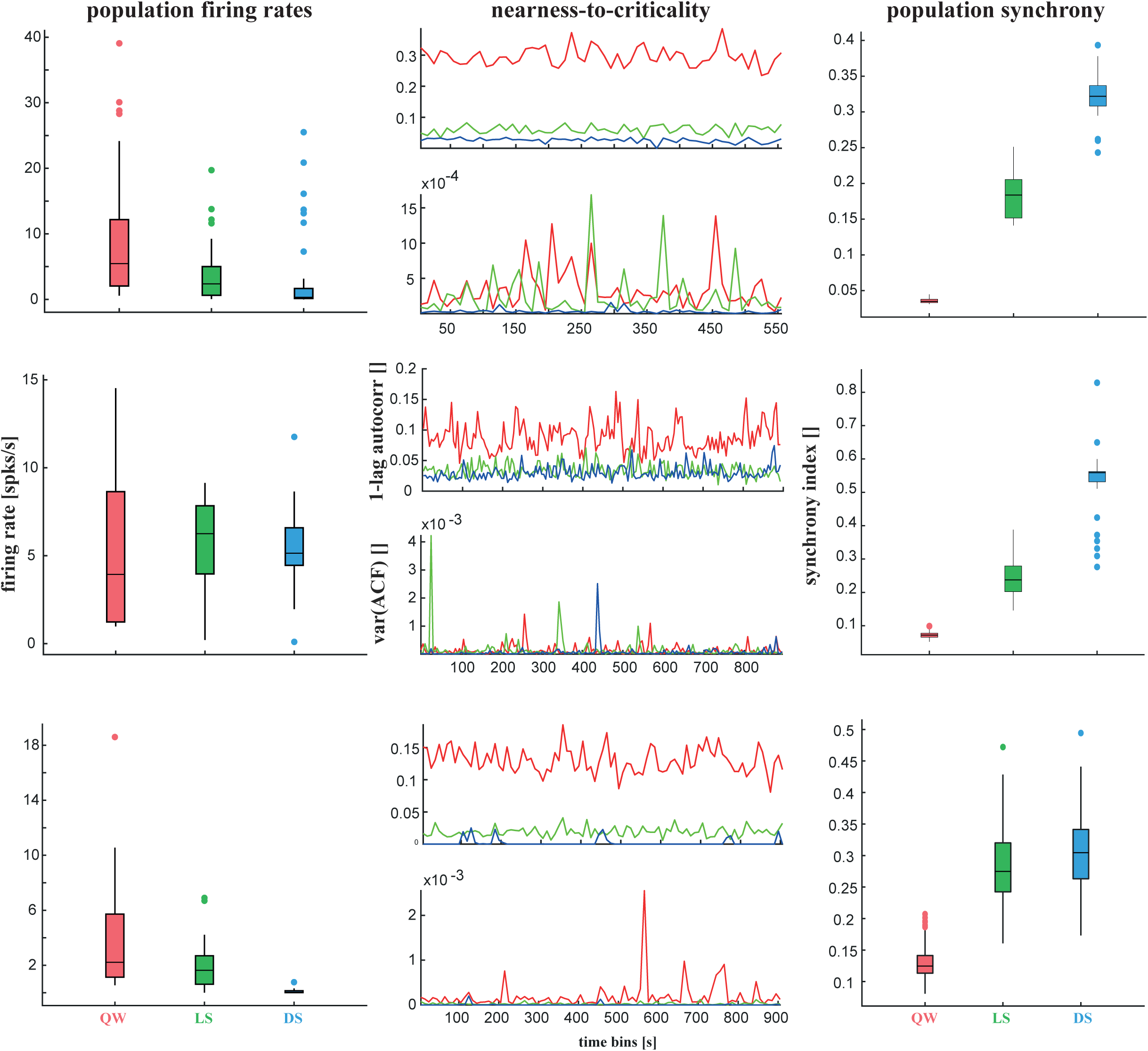
Population characteristics under different states. Top row - Col 1 - A comparison of firing rates of the fixed population (based on the firing rates during QW, fr >= 0.5) during different states of (un) consciousness in Monkey A. Col 2 - Metrics of nearness-to-criticality and critical slowing, in 10s bins during different states of (un)consciousness. Col 3 - A comparison of population synchrony of the fixed population of neurons, during different states of (un)consciousness (PFC) Middle Row - Same as above, for Monkey B (PFC) Bottom Row - Same as above, for Monkey C (PPC)

**SI Figure 2.**
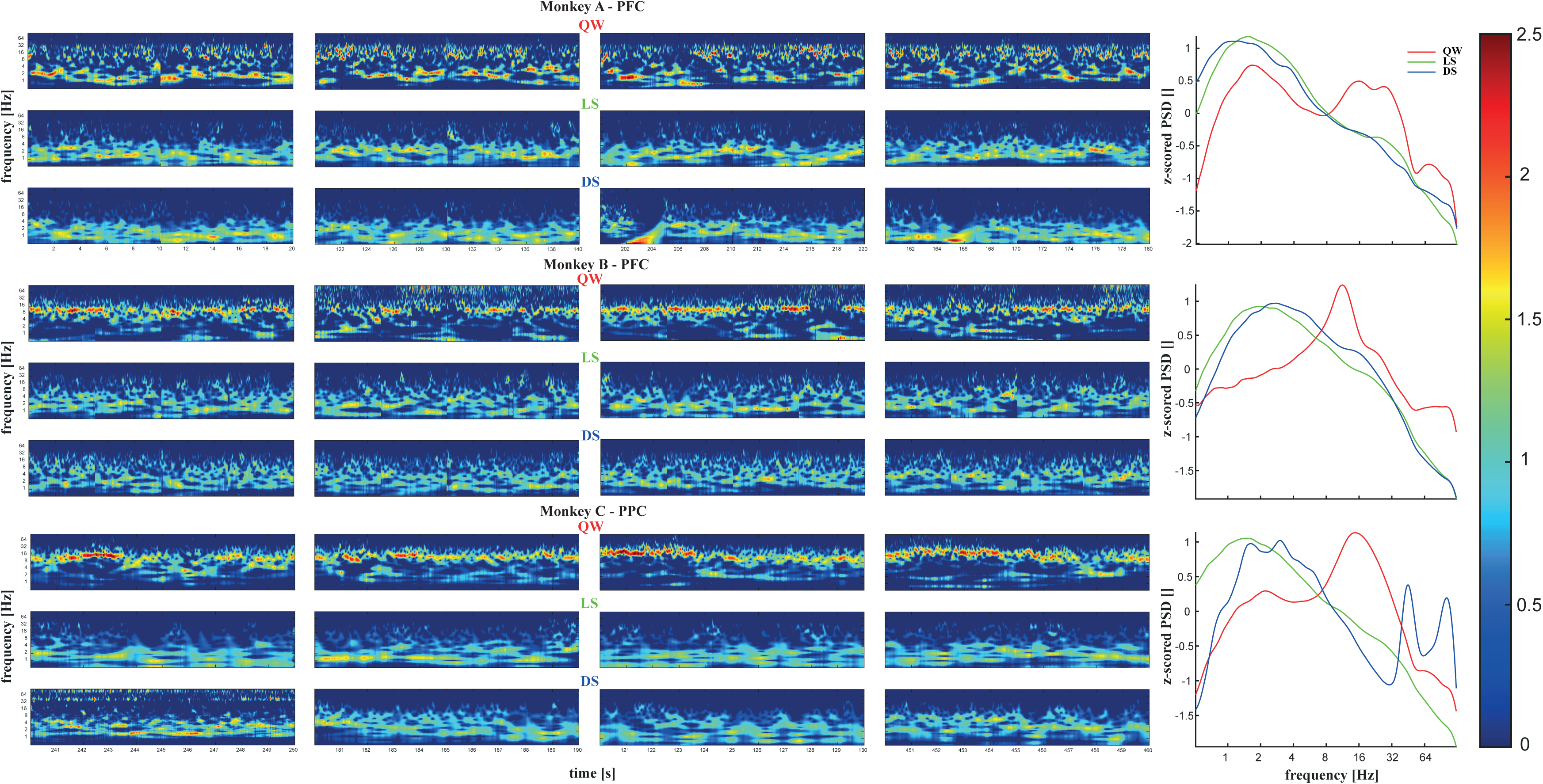
Examples of QW, LS and DS. More examples of QW, LS and DS periods, from PFC and PPC Top rows in each subplot correspond to QW, middle to LS and lower to DS. Sideplots show the comparison of the collapsed power spectral density (z-scored). The two peaks at 50 and 100Hz are non-physiological, and come from strong line noise during this particular epoch of this particular recording session.

**SI Figure 3.**
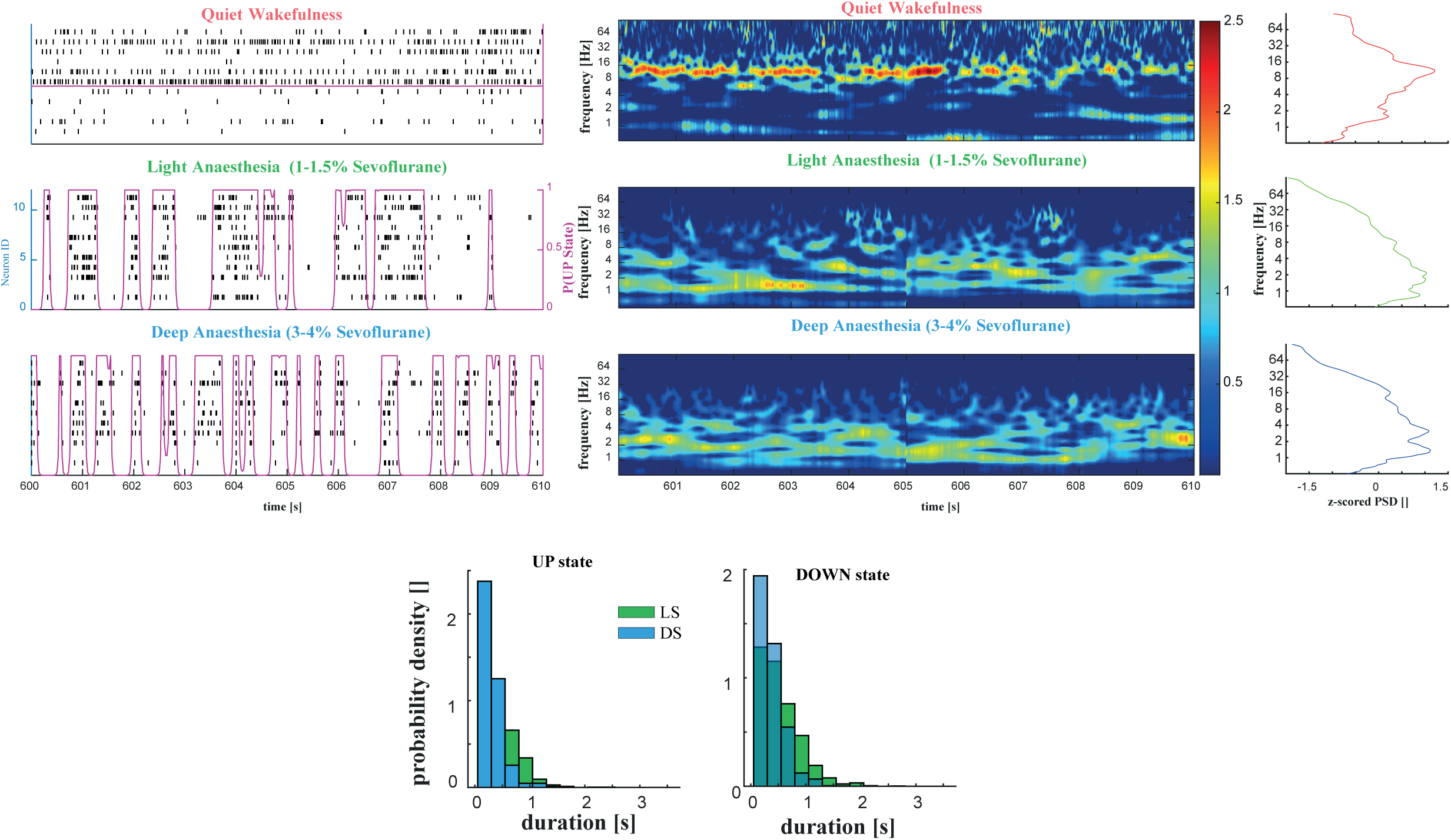
Resting state characteristics for Monkey B - PFC. The raster plot of the fixed population displays sparse and desynchronised spiking during QW (top row), while it displays the characteristic UP/DOWN states during both states of anaesthesia. The time-frequency decomposition and the the power spectral density both show significantly stronger low-frequency activity during LS and DS, but higher beta and gamma activity during QW. Although DS was maintained at 3%, there was no significant lengthening of the silent periods as compared to LS (1%). Additionally, there were no significant differences observed between the durations of the UP and DOWN states, pointing to no difference in anaesthesia-induced cortical entrainment.

**SI Figure 4.**
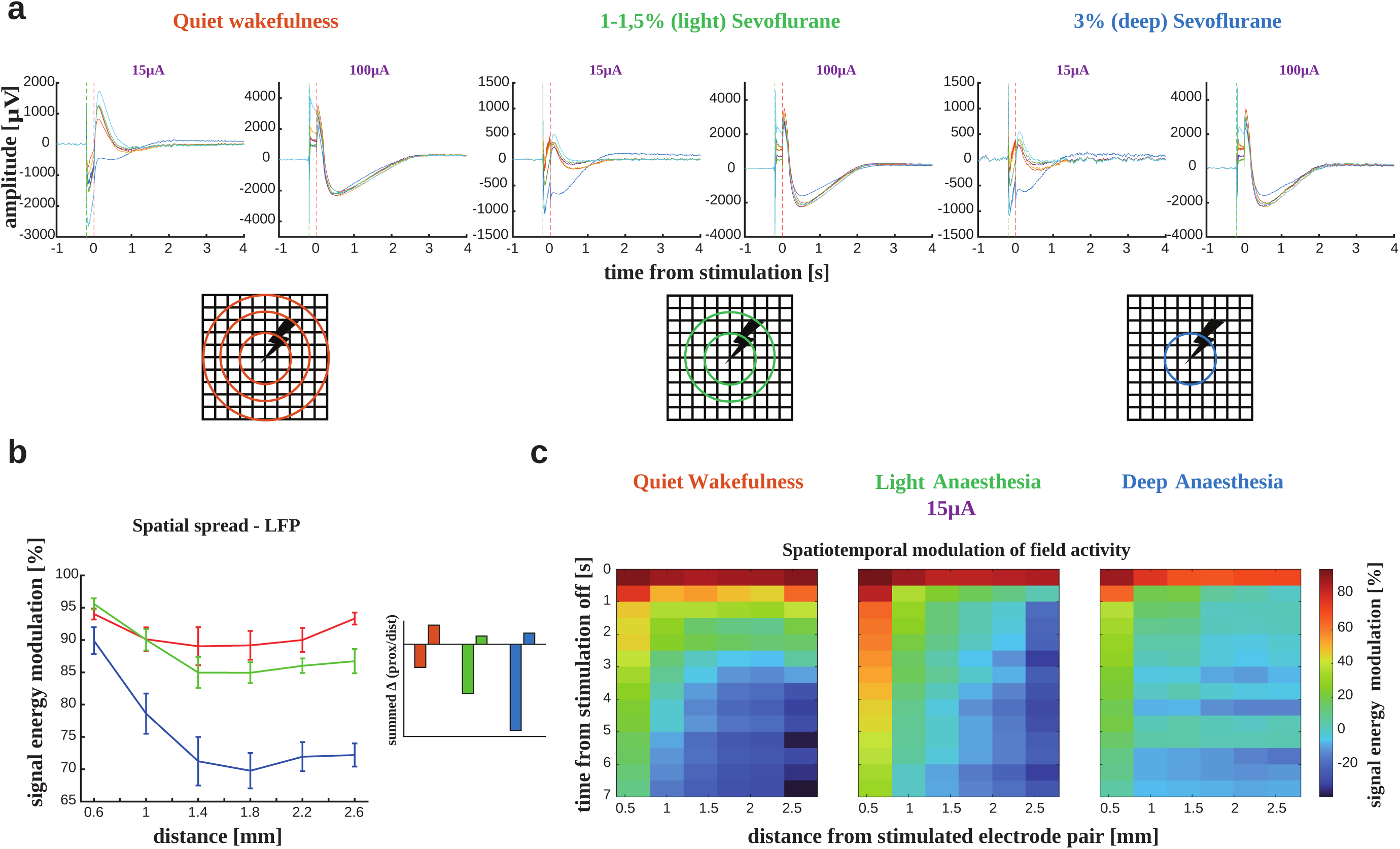
Spatial and temporal patterns of modulation for Monkey B - PFC. A - Please refer to Fig. 5 in the main text for a description B - While a conclusive non-monotonic pattern is observed in the modulation in the LFPs in QW, the strength of modulation falls of linearly during both LS and DS in LFPs. Although the sum of rate of change in modulation in all three states do display a weak distal increase, it is not established to be conclusive statistically. C - Spatio-temporal modulation of field activity.

1 Table for Amplitude: 1 μA
| State | Transient Timing [s] | Dispersal | Transient Strength [corrSpks/s] | Dispersion | Rebound Timing [s] | Dispersal | Rebound Strength [corrSpks/s] | Dispersal |
| --- | --- | --- | --- | --- | --- | --- | --- | --- |
| QW | 0.8438 ± 0.0039 | 0.6562 | 93.3845 ± 0.3070 | 3974.9238 | n/a | n/a | n/a | n/a |
| LS | 1.3086 ± 0.0047 | 1.3967 | 26.7084 ± 0.0812 | 419.2467 | n/a | n/a | n/a | n/a |
| DS | 0.7064 ± 0.0043 | 0.5723 | 206.2554 ± 0.7104 | 15490.6126 | n/a | n/a | n/a | n/a |

4 CIs for Amplitude: 1 μA
|  | QW vs LS | LS vs DS | QW vs DS |
| --- | --- | --- | --- |
| Time | CIs:0.0003;0.0162 | CIs:0.0003;0.0161 | CIs:0.0003;0.0156 |
| Amplitude | CIs:0.0076;0.6921 | CIs:0.0151;1.9451 | CIs:0.0407;1.7713 |

6 Table for Amplitude: 5 μA
| State | Transient Timing [s] | Dispersal | Transient Strength [corrSpks/s] | Dispersion | Rebound Timing [s] | Dispersal | Rebound Strength [corrSpks/s] | Dispersal |
| --- | --- | --- | --- | --- | --- | --- | --- | --- |
| QW | 0.7572 ± 0.0026 | 0.4634 | 103.1004 ± 0.2183 | 3266.2309 | n/a | n/a | n/a | n/a |
| LS | 1.1009 ± 0.0032 | 1.0049 | 39.4883 ± 0.0889 | 756.4541 | n/a | n/a | n/a | n/a |
| DS | 0.5687 ± 0.0024 | 0.3310 | 262.2674 ± 0.6476 | 23749.6579 | n/a | n/a | n/a | n/a |

8 CIs for Amplitude: 5 μA
|  | QW vs LS | LS vs DS | QW vs DS |
| --- | --- | --- | --- |
| Time | CIs:0.0001;0.0091 | CIs:0.0001;0.0113 | CIs:0.0001;0.0067 |
| Amplitude | CIs:0.0193;0.4975 | CIs:0.046;1.5001 | CIs:0.012;1.6153 |

11 Table for Amplitude: 8 μA
| State | Transient Timing [s] | Dispersal | Transient Strength [corrSpks/s] | Dispersal | Rebound Timing [s] | Dispersal | Rebound Strength [corrSpks/s] | Dispersal |
| --- | --- | --- | --- | --- | --- | --- | --- | --- |
| QW | 0.7023 ± 0.0022 | 0.4075 | 102.6141 ± 0.2063 | 3492.3023 | n/a | n/a | n/a | n/a |
| LS | 1.1668 ± 0.0028 | 0.7758 | 32.5500 ± 0.0682 | 471.1069 | n/a | n/a | n/a | n/a |
| DS | 0.5629 ± 0.0019 | 0.2651 | 178.6599 ± 0.3636 | 10156.9429 | n/a | n/a | n/a | n/a |

13 CIs for Amplitude: 8 μA
|  | QW vs LS | LS vs DS | QW vs DS |
| --- | --- | --- | --- |
| Time | CIs:0.0002;0.0071 | CIs:0.0001;0.0086 | CIs:0.0002;0.006 |
| Amplitude | CIs:0.0117;0.4993 | CIs:0.0152;0.976 | CIs:0.0074;0.8898 |

15 Table for Amplitude: 15 μA
| State | Transient Timing [s] | Dispersal | Transient Strength [corrSpks/s] | Dispersal | Rebound Timing [s] | Dispersal | Rebound Strength [corrSpks/s] | Dispersal |
| --- | --- | --- | --- | --- | --- | --- | --- | --- |
| QW | -0.0739 ± 0.0015 | 0.0234 | 64.3858 ± 0.2950 | 967.5470 | 1.8880 ± 0.0032 | 0.3152 | 26.6120 ± 0.1254 | 473.0090 |
| LS | 0.1379 ± 0.0025 | 0.1121 | 25.1672 ± 0.1277 | 281.6930 | 2.0960 ± 0.0033 | 0.6020 | 17.4788 ± 0.0598 | 192.9809 |
| DS | -0.1092 ± 0.0005 | 0.0028 | 57.5013 ± 0.2630 | 755.9151 | 1.5646 ± 0.0038 | 0.4476 | 41.5794 ± 0.2023 | 1260.3483 |

17 CIs for Amplitude: 15 μA
|  | QW vs LS | LS vs DS | QW vs DS | Rebound QW vs LS | Rebound LS vs DS | Rebound QW vs DS |
| --- | --- | --- | --- | --- | --- | --- |
| Time | CIs:0.0001;0.0077 | CIs:0.0002;0.0076 | CIs:0;0.0035 | CIs:0.0002;0.0111 | CIs:0.0002;0.0126 | CIs:0.0002;0.0125 |
| Amplitude | CIs:0.0126;<br>0.8382 | CIs:0.0097;<br>0.7864 | CIs:0.0107;<br>0.9128 | CIs:0.0044;<br>0.2845 | CIs:0.0068;<br>0.4393 | CIs:0.0073;<br>0.5404 |

20 Table for Amplitude: 20 μA
| State | Transient Timing [s] | Dispersal | Transient Strength [corrSpks/s] | Dispersal | Rebound Timing [s] | Dispersal | Rebound Strength [corrSpks/s] | Dispersal |
| --- | --- | --- | --- | --- | --- | --- | --- | --- |
| QW | 0.0784 ± 0.0019 | 0.0450 | 43.3549 ± 0.3318 | 1336.7909 | 2.1949 ± 0.0031 | 0.2705 | 22.3349 ± 0.1030 | 299.4019 |
| LS | 0.1974 ± 0.0017 | 0.0852 | 31.8037 ± 0.1786 | 963.2913 | 2.4295 ± 0.0030 | 0.3984 | 13.3161 ± 0.0519 | 116.8168 |
| DS | 1.3361 ± 0.0050 | 0.7478 | 68.4755 ± 0.2659 | 2098.2474 | 3.0582 ± 0.0096 | 0.3002 | 95.1274 ± 1.4681 | 7091.3451 |

22 CIs for Amplitude: 20 μA
|  | QW vs LS | LS vs DS | QW vs DS | Rebound QW vs LS | Rebound LS vs DS | Rebound QW vs DS |
| --- | --- | --- | --- | --- | --- | --- |
| Time | CIs:0.0001; 0.007 | CIs:0.0002; 0.016 | CIs:0.0003; 0.0217 | CIs:0.0002; 0.0104 | CIs:0.0004; 0.0263 | CIs:0.0002; 0.0224 |
| Amplitude | CIs:0.0108; 0.8304 | CIs:0.0113; 0.7856 | CIs:0.0136; 1.0292 | CIs:0.004; 0.2554 | CIs:0.0154; 1.3314 | CIs:0.0277; 1.5633 |

24 Table for Amplitude: 25 μA
| State | Transient Timing [s] | Dispersal | Transient Strength [corrSpks/s] | Dispersal | Rebound Timing [s] | Dispersal | Rebound Strength [corrSpks/s] | Dispersal |
| --- | --- | --- | --- | --- | --- | --- | --- | --- |
| QW | 0.0803 ± 0.0015 | 0.0323 | 31.4526 ± 0.2416 | 799.9227 | 2.1955 ± 0.0031 | 0.2691 | 21.2145 ± 0.1012 | 285.4046 |
| LS | 0.2491 ± 0.0018 | 0.0959 | 34.4576 ± 0.1941 | 1162.2963 | 2.5425 ± 0.0030 | 0.3828 | 15.7265 ± 0.0624 | 161.8122 |
| DS | 0.0693 ± 0.0014 | 0.0183 | 49.5116 ± 0.4471 | 1889.1332 | 2.1147 ± 0.0024 | 0.1778 | 67.1164 ± 0.2642 | 2212.6116 |

26 CIs for Amplitude: 25 μA
|  | QW vs LS | LS vs DS | QW vs DS | Rebound | Rebound | Rebound |
| --- | --- | --- | --- | --- | --- | --- |
|  |  |  |  | QW vs LS | LS vs DS | QW vs DS |
| Time | CIs:0.0001;<br>0.0062 | CIs:0.0001;<br>0.0076 | CIs:0.0001;<br>0.0049 | CIs:0.0002;<br>0.0107 | CIs:0.0002;<br>0.0098 | CIs:0.0001;<br>0.0081 |
| Amplitude | CIs:0.0107;<br>0.7699 | CIs:0.012;0<br>.9709 | CIs:0.0118;<br>1.0996 | CIs:0.0041;<br>0.2523 | CIs:0.012;0.<br>6456 | CIs:0.0069;<br>0.8019 |

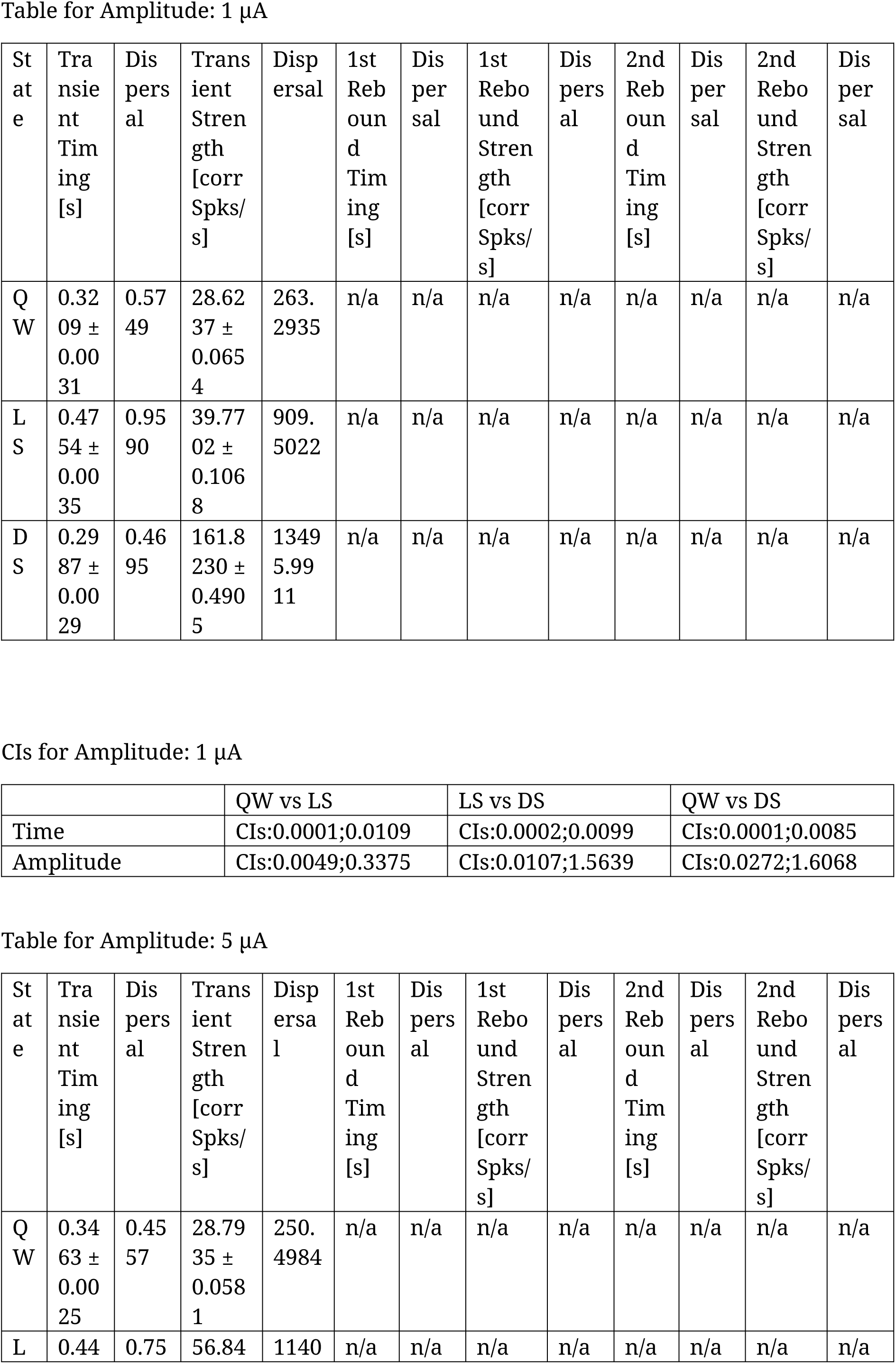
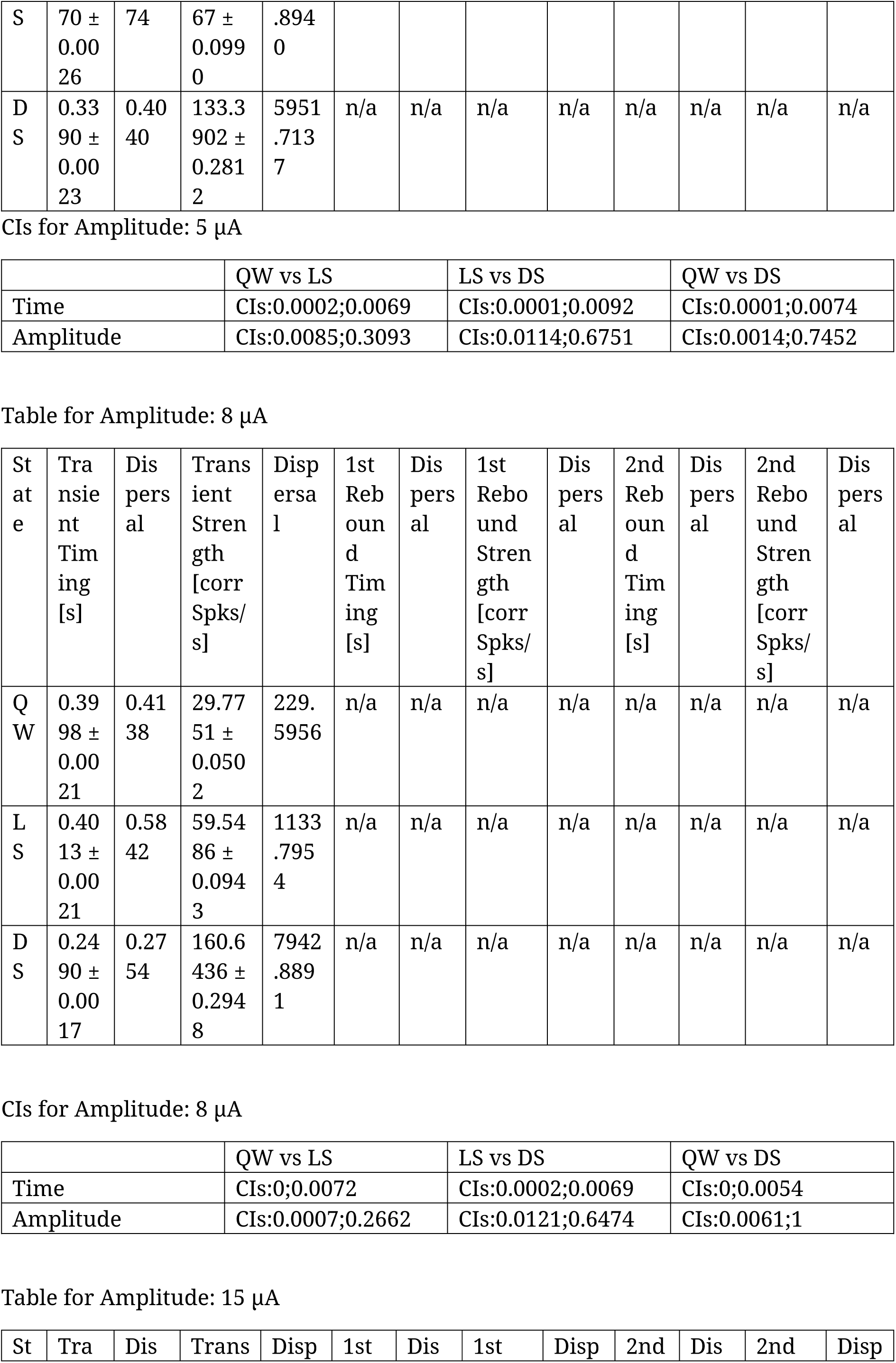
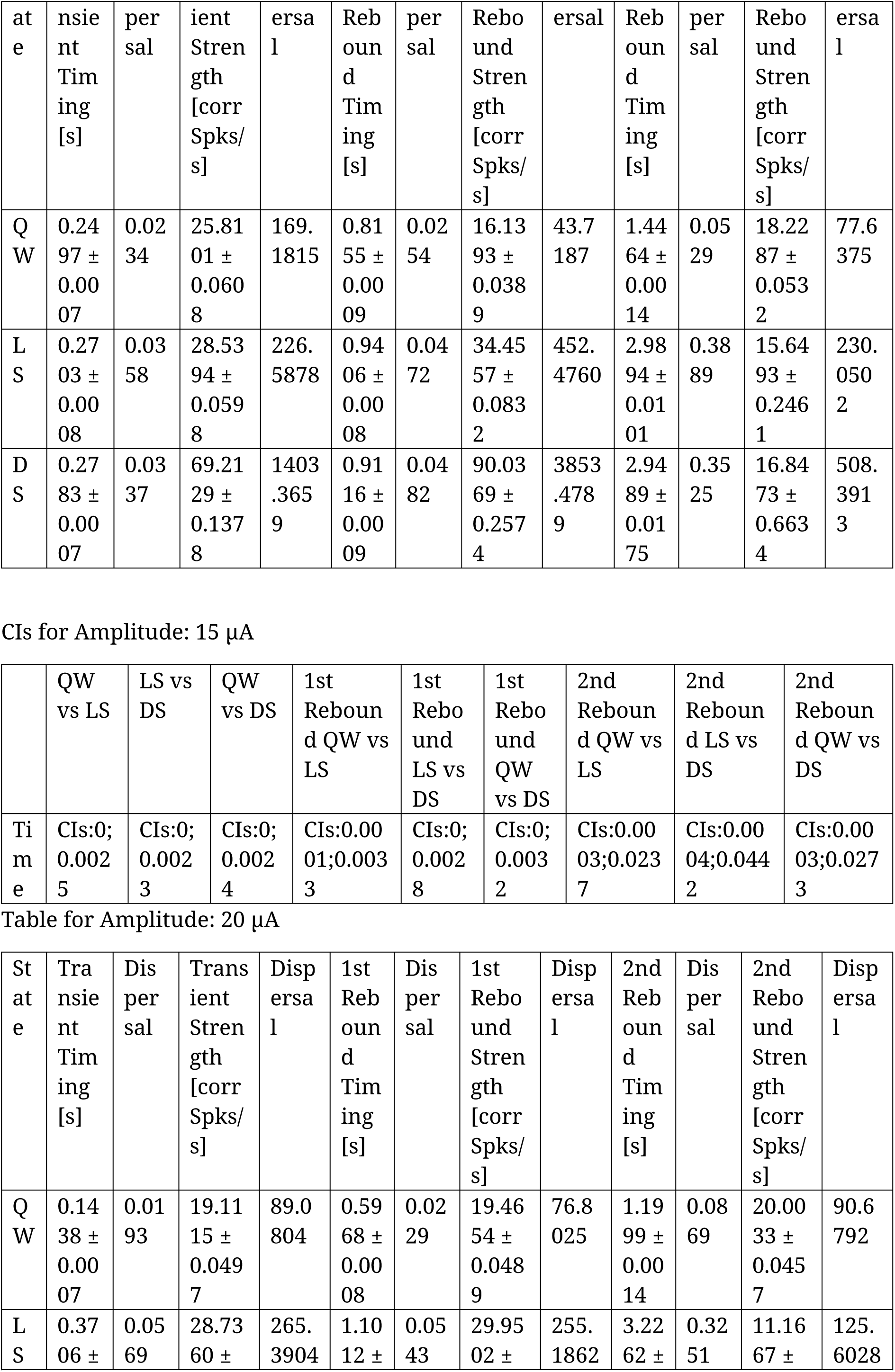
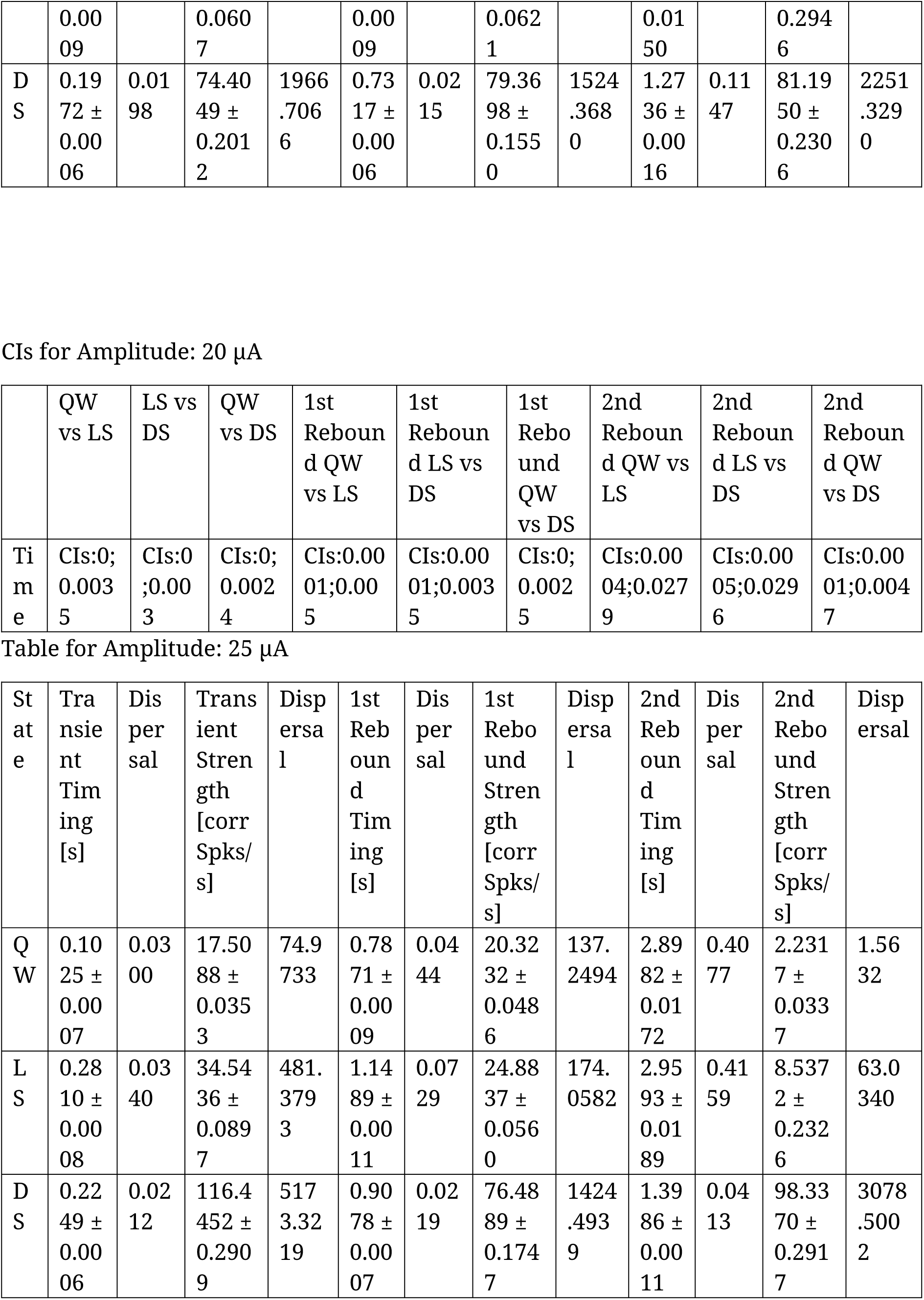
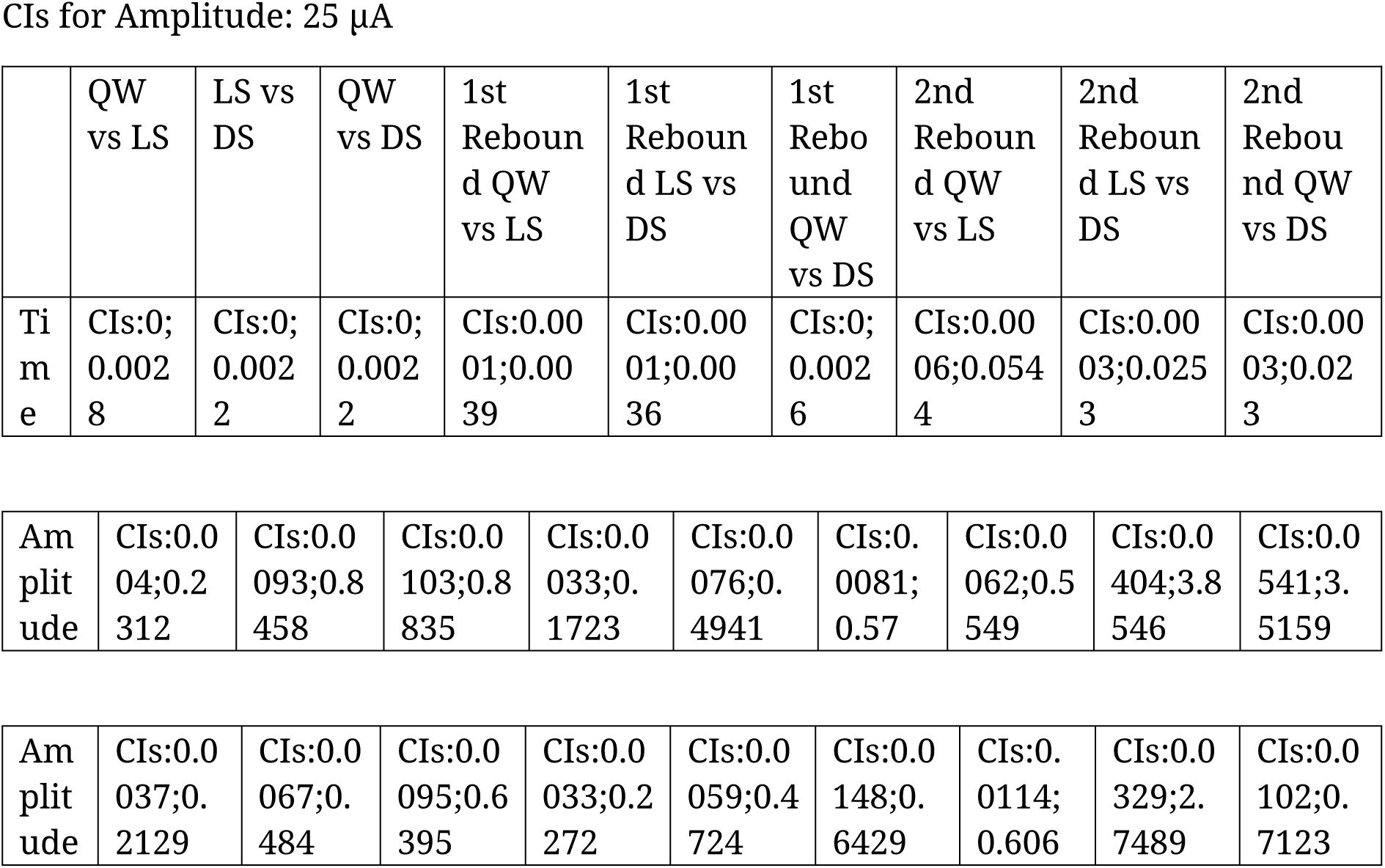
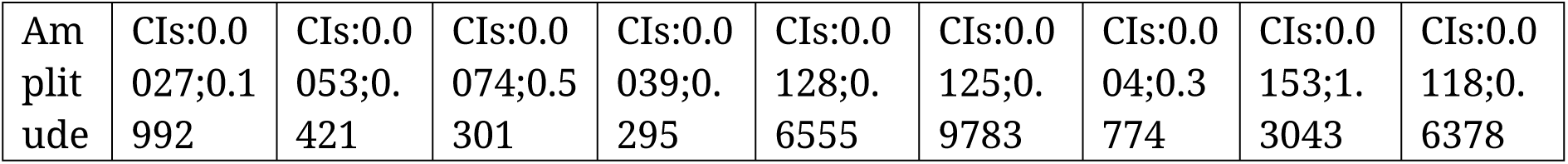

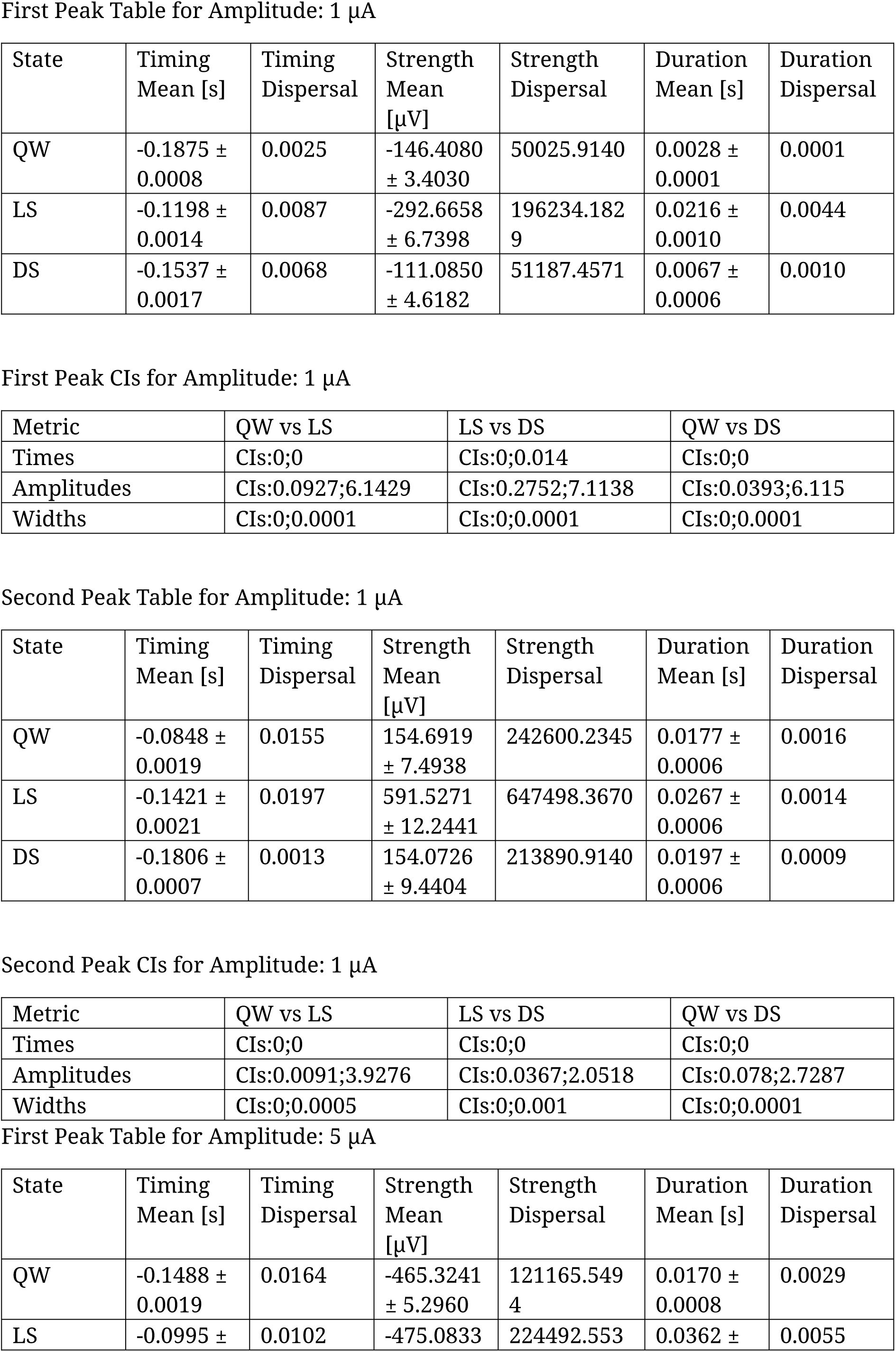
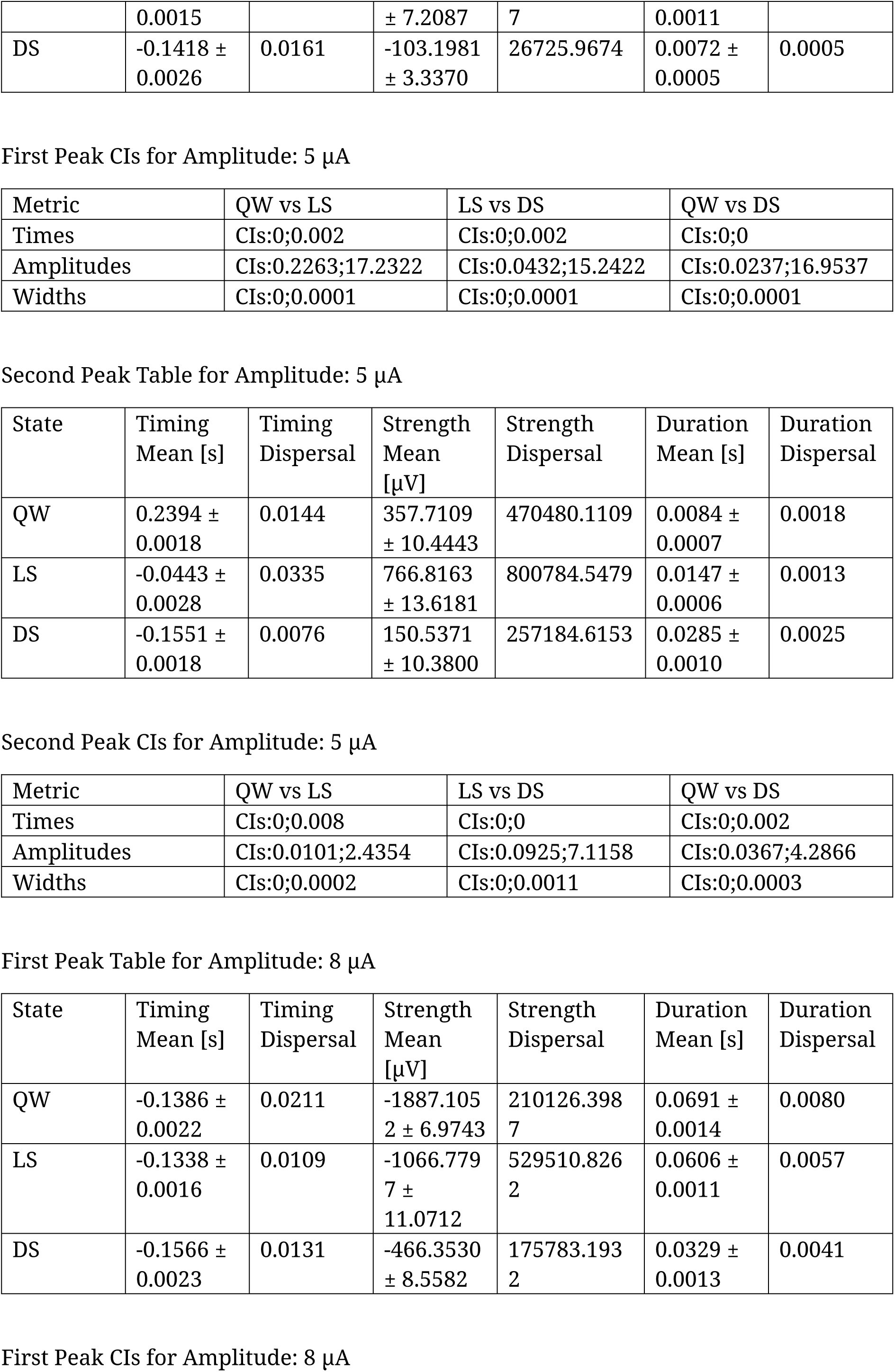
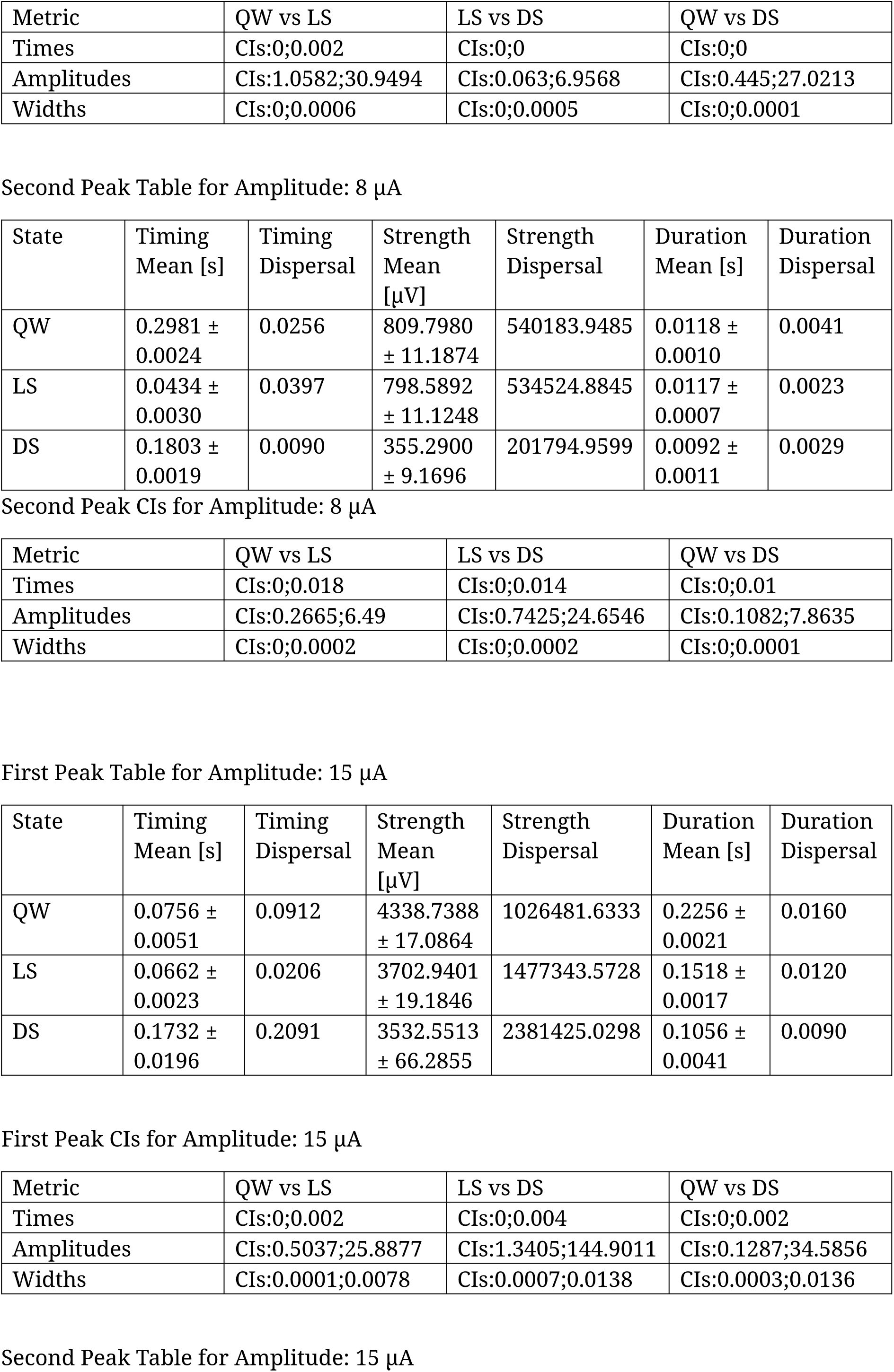
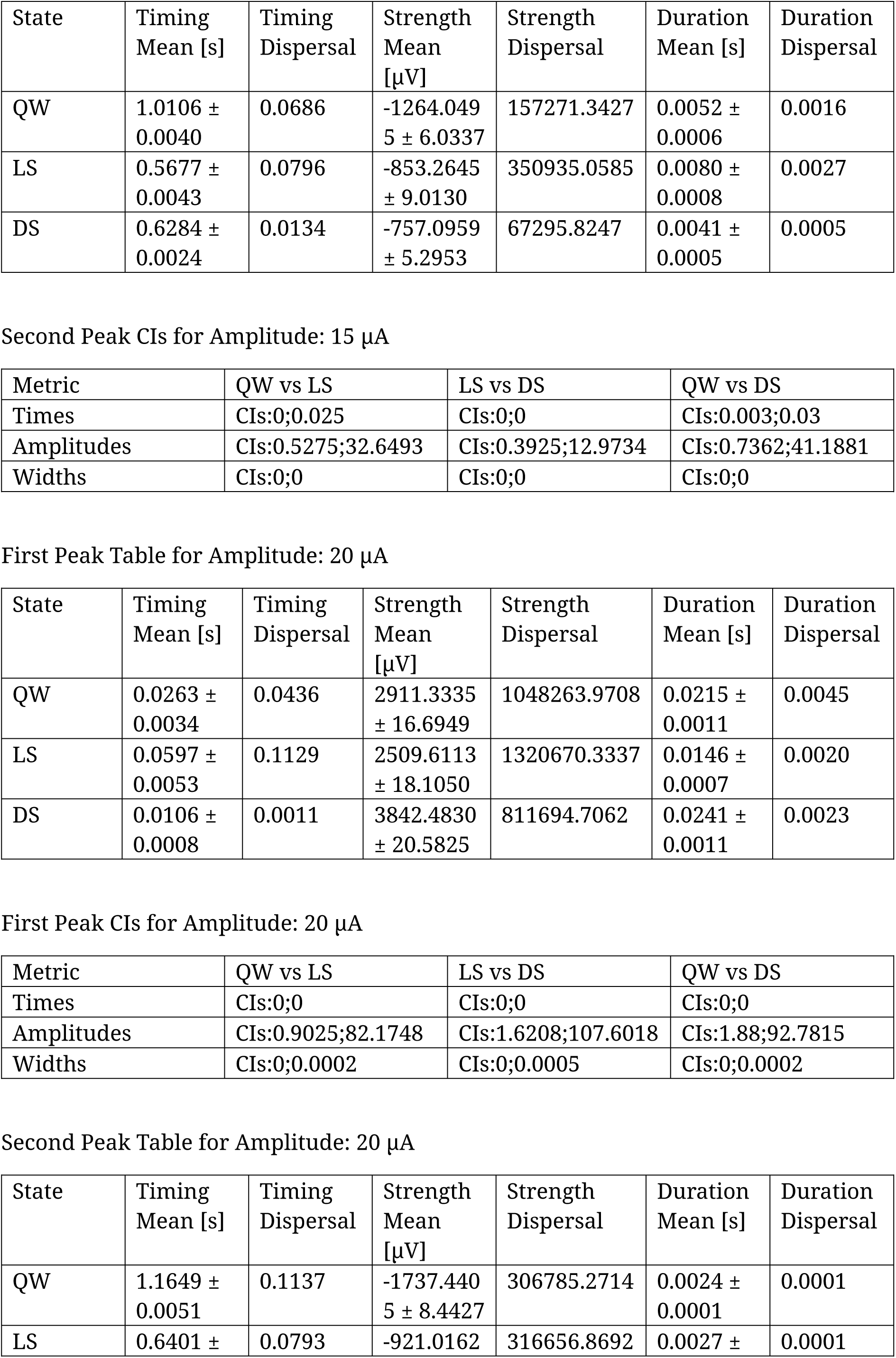
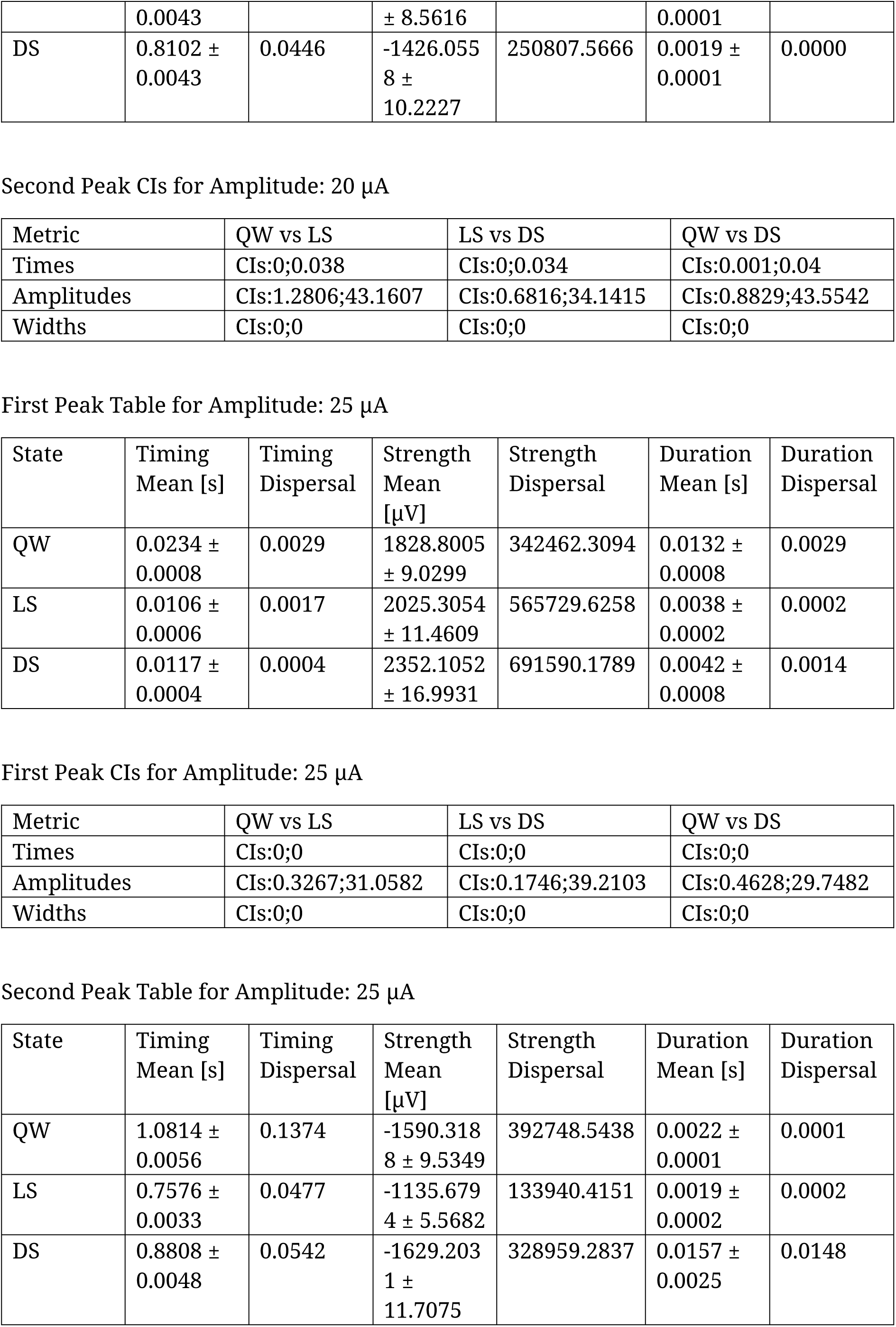
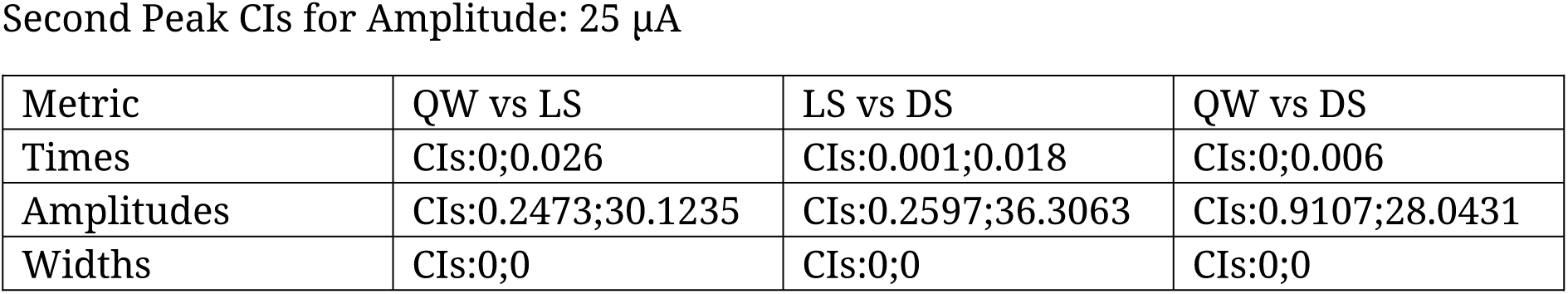

## Notes

### Competing Interest Statement

The authors have declared no competing interest.

### Summary of Updates

Changed title Changed author list Changed introduction Other minor changes

